# ARMH3 acts as a central scaffold at the Golgi/TGN through interactions with Arl5, GBF1, and PI4KB

**DOI:** 10.64898/2026.04.15.718671

**Authors:** Mackenzie K Scott, Giselle CT Klynsoon, Emma E Walsh, Sushant Suresh, Hunter G Nyvall, John E Burke

**Author notes:** Lead contact: John E Burke.

## Abstract

The armadillo repeat protein ARMH3 regulates the activity and localisation of the Golgi resident lipid kinase phosphatidylinositol 4 kinase IIIβ (PI4KB) and the Golgi-specific brefeldin A-resistance guanine nucleotide exchange factor 1 (GBF1) that activates Arf1. ARMH3 localises to the trans Golgi network (TGN) via the GTPase Arl5. We used hydrogen deuterium exchange mass spectrometry (HDX-MS) and AI-enabled modeling to define the interfaces of ARMH3 with its binding partners Arl5, PI4KB, and GBF1. The ARMH3-Arl5 interface was determined to consist of the N and C termini of ARMH3, with Arl5 binding causing conformational changes in ARMH3 located at a shared PI4KB/GBF1 interface. Both PI4KB and GBF1 form mutually exclusive complexes with ARMH3, with GBF1 binding to ARMH3 through a disordered loop we have named the ARMH3 binding region (ABR). The ARMH3 interfaces in PI4KB and GBF1 contain phosphosites, with the phosphomimetic mutation of GBF1 blocking complex formation. These findings provide new insights into the role of ARMH3 as a master coordinator of GTPase and phosphoinositide signaling at the Golgi/TGN.

## Introduction

The Golgi apparatus is the central sorting hub of the secretory pathway, directing the trafficking of newly synthesized proteins and lipids to their appropriate cellular destinations (Glick & Nakano, 2009). The proper functioning and formation of the Golgi is coordinated by both lipid phosphoinositides and members of the Ras superfamily of small GTPases. Two of the more well studied factors that play critical roles in the Golgi are the phosphoinositide species phosphatidylinositol 4 phosphate (PI4P) and the ADP-ribosylation factor GTPase Arf1, which together play important roles in cargo selection and lipid transfer that are necessary for Golgi maintenance and trafficking (Taylor *et al*, 2025; De Matteis *et al*, 2025; Wong *et al*, 2019). Phosphoinositide production is controlled by the action of phosphoinositide kinases and phosphatases (Burke, 2018; Dyson *et al*, 2012) that mediate transfer of phosphate to the phosphatidylinositol headgroup. Arf1 is a GTPase that cycles between an inactive guanosine diphosphate (GDP)-bound state and an active guanosine triphosphate (GTP)-bound state, with guanine nucleotide exchange factors mediating GDP to GTP exchange (Nawrotek *et al*, 2016). Defining the molecular mechanisms that control how activation of the lipid kinases and guanine nucleotide exchange factors (GEFs) that generate PI4P and activate Arf GTPases is essential to define how Golgi signalling is regulated.

PI4P is found throughout the different regions of the Golgi, but its levels are higher in the Trans-Golgi and Trans-Golgi-Network (TGN). PI4P at the Golgi and TGN are produced by the lipid kinases phosphatidylinositol 4-kinase IIIβ (PI4KB) and PI4KIIα (PI4K2A) (Boura & Nencka, 2015; Dornan *et al*, 2016). PI4KB is a cytosolic enzyme that is recruited to the Golgi/TGN through its interactions with multiple scaffolding proteins, including the Golgi protein Acyl CoA binding protein 3 (ACBD3) (Klima *et al*, 2016; McPhail *et al*, 2017) and the TGN localised protein Armadillo-like helical domain-containing protein 3 (ARMH3, also known as C10orf76) (McPhail *et al*, 2020; Ishida *et al*, 2024; Fang *et al*, 2023; Mizuike *et al*, 2023). Its recruitment by separate Golgi/TGN scaffolds allows for distinct PI4KB signalling pools that can have non-overlapping functions (Mizuike *et al*, 2023). Arf1 plays an important role in controlling the activity of PI4KB through an unknown molecular mechanism (Godi *et al*, 1999), with Arf1 activation at the Golgi being controlled by the action of Golgi-specific brefeldin A-resistance guanine nucleotide exchange factor 1 (GBF1) that mediates the exchange of GDP to GTP in Arf1. ARMH3 is also a known regulator of GBF1, with its knockout leading to Golgi disruption and decreased GBF1 recruitment to the Golgi (Chan *et al*, 2019). Together these studies have identified ARMH3 as a critical regulator of both PI4P and Arf1 levels at the Golgi and TGN through its interactions with PI4KB and GBF1, however, the full molecular details for how this occurs is unknown.

GBF1 is a large multidomain protein, with Golgi recruitment regulated by many additional factors beyond ARMH3, including an amphipathic membrane binding helix (Bouvet *et al*, 2013; Muccini *et al*, 2022), binding to specific phosphoinositides (Meissner *et al*, 2018), phosphorylation (Walton *et al*, 2023), and interaction with GTP-loaded Rab1 (Monetta *et al*, 2007). The cryo EM structure of the yeast paralog GEA1 revealed that it is a large dimeric complex composed of a DCB-HUS domain (dimerization and cyclophilin binding, homology upstream of Sec7), a Sec7 GEF domain, and three HDS domains (homology downstream) of Sec7 (Muccini *et al*, 2022). GEA1 is able to adopt unique open and closed forms, with nucleotide free Arf1 binding stabilising the open state. Regulators of GBF1 at the Golgi could therefore be predicted to either alter membrane recruitment or conformation at the Golgi. While no high resolution structural information exists for human GBF1, AlphaFold modeling (Varadi *et al*, 2022) suggest that it is structurally highly similar. Differential proteolytic digestion between cytosolic human GBF1 and membrane bound GBF1 suggests that it may also cycle between open and closed forms (Meissner *et al*, 2023). Defining how different binding partners and post-translational modifications alter the conformation of GBF1 is critical for understanding its regulation.

The role of ARMH3 in regulating trafficking at the Golgi/TGN has only recently been established, with it being identified as an essential gene that regulates TGN PI4P levels (Blomen *et al*, 2015). ARMH3 is recruited to the TGN by GTP-loaded Arl5, which is a member of the Arf family of GTPases (Ishida *et al*, 2024). Arl5 in turn is recruited to the TGN by its interactions with the SYS1-ARFRP1 complex. This SYS1-ARFRP1-ARL5-ARMH3-PI4KB pathway is critical for generating a TGN PI4P pool, which is required for the correct localisation and stability of glycosylation enzymes. However, how ARMH3 interacts with Arl5, and how this may alter the interactions with PI4KB or GBF1 is unknown. The binding site of PI4KB to ARMH3 is located at a disordered linker in the N-lobe of the PI4KB kinase domain, which interacts with a region near the C-terminus of ARMH3 (McPhail *et al*, 2020). This ARMH3 binding site in PI4KB can be phosphorylated by protein kinase A (PKA) (Isobe *et al*, 2017), with phosphorylation leading to decreased binding to ARMH3 (McPhail *et al*, 2020). Overall, this suggests that ARMH3 can make myriad interactions with a variety of protein binding partners, and that these associations can be regulated by post-translational modifications (PTMs).

We hypothesized that the function of ARMH3 is dependent on its ability to form complexes with Arl5, PI4KB, and GBF1. To define the molecular basis for how ARMH3 is able to interact with these proteins we utilised a synergy of hydrogen deuterium exchange mass spectrometry (HDX-MS) and AlphaFold3 molecular modeling. This allowed for the design of complex disrupting mutations that both validated the binding sites, and can be used in future *in vivo* analysis of ARMH3. We find that Arl5 binds to a site at the N and C termini of ARMH3, with this interaction causing extensive allosteric conformational changes. The Arl5 site is distinct from a shared PI4KB and GBF1 binding interface on ARMH3. The interfaces of PI4KB and GBF1 with ARMH3 both contain phosphosites that modulate ARMH3 binding. Together, these findings provide a mechanistic framework for understanding how ARMH3 functions as a central scaffolding node that coordinates Arf activation and PI4P synthesis at the TGN, with direct implications for understanding its roles in both normal membrane trafficking and its subversion during disease.

## Results

To investigate the molecular basis of the Arl5-ARMH3 interaction, we produced recombinant forms of both proteins. Like other Arf family GTPases, Arl5 contains an N-terminal amphipathic helix that is myristoylated at G2, which mediates nucleotide loading and enables membrane association (Wang *et al*, 2005). To ensure proper nucleotide loading, we generated N-terminal truncations of both Arl5A and B beginning at Q15, consistent with previous studies (Ishida *et al*, 2024), which are referred to as Arl5A and Arl5B throughout this paper (**Fig 1A**). We generated full-length ARMH3 in both *Spodoptera frugiperda* (*Sf9*) cells as previously described (McPhail *et al*, 2020), as well as in *Escherichia coli,* which displayed a similar molecular weight and gel filtration profile to the *Sf9*-derived construct. To quantify the nucleotide-dependent GTPase-effector interaction of ARMH3 with Arl5 (Ishida *et al*, 2024), we carried out biolayer interferometry (BLI) assays using His-tagged Arl5A(GTPγS), Arl5B(GTPγS), and Arl5B(GDP) immobilised on the biosensor tip and ARMH3 as the analyte. ARMH3 selectively bound the GTPγS-loaded form of both Arl5A and Arl5B (**Fig 1B+C)**. To measure the kinetics of the ARMH3-Arl5B(GTPγS) interaction, immobilised His-tagged Arl5B(GTPγS) (25 nM) was dipped into varying concentrations (15-100 nM) of ARMH3. The dissociation constant of the Arl5B(GTPγS)-ARMH3 complex was 75.38 ± 18.04 nM, calculated as the mean K_D_ across all curves meeting the inclusion criteria (**Fig 1D**). Average binding parameters (K_D_, K_on_, K_off_) are shown in Table S1, raw data and binding parameters for each individual curve are available in the source data.

**Figure 1:**
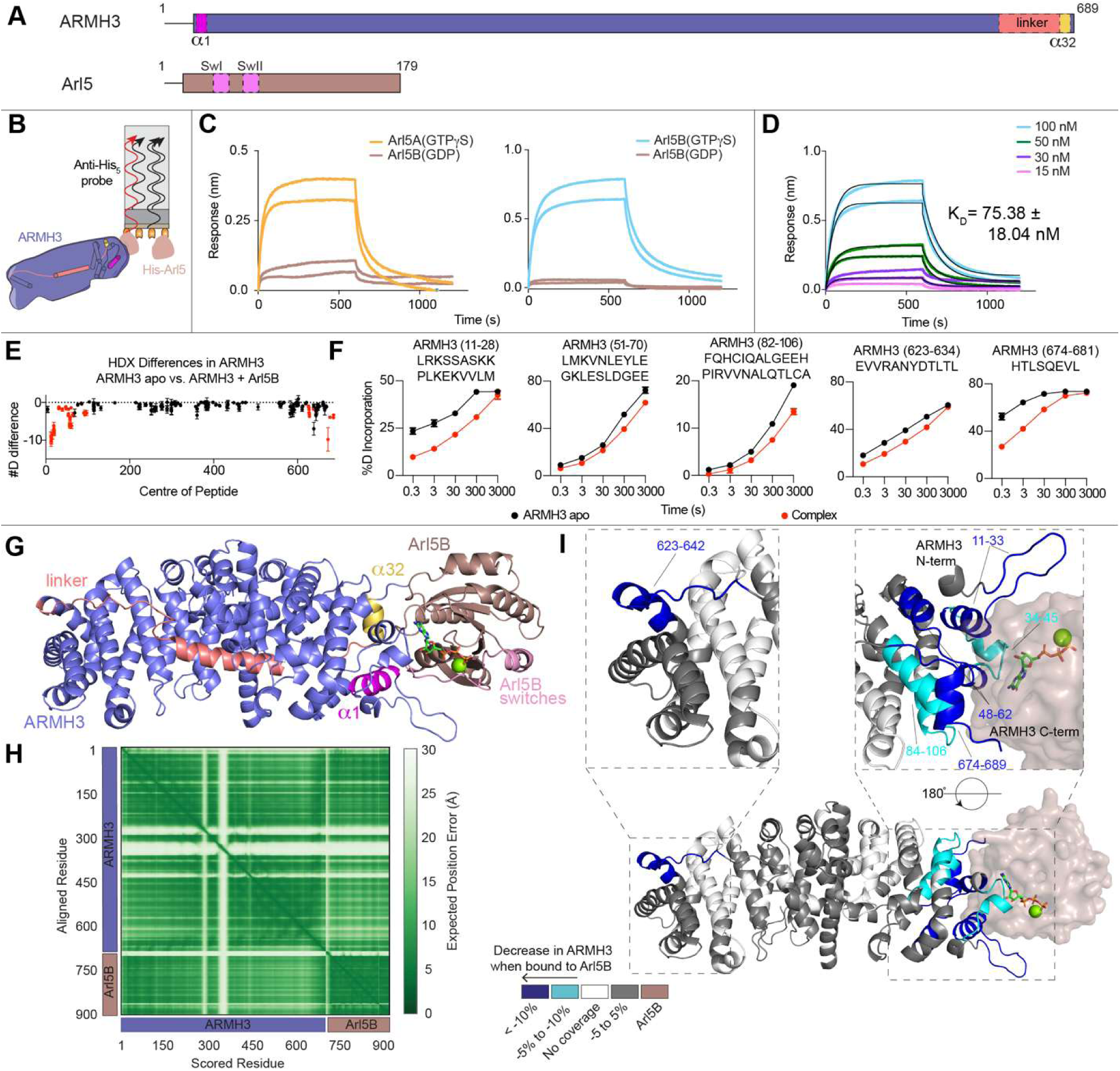
Biolayer interferometry and HDX-MS reveal canonical GTPase-effector interaction. **A)** Domain schematics of full-length ARMH3 and Arl5. Constructs used in this paper are Arl5A(15-179) and Arl5B(15-179), which are referred to as Arl5A and Arl5B throughout this paper. **B)** Schematic of the biolayer interferometry (BLI) assay showing binding of immobilised His-Arl5 on the tip to full-length ARMH3 in solution. **C)** BLI traces of Arl5A(GTPγS), Arl5B(GTPγS), and Arl5B(GDP) binding to full-length ARMH3. His-Arl5(GTPγS) or His-Arl5B(GDP) was loaded onto the anti-penta His tip at 25 nM and dipped into ARMH3 at 100 nM. **D)** Dose response of Arl5B(GTPγS) binding to ARMH3. His-Arl5B(GTPγS) was loaded onto the anti-penta His tip at 25 nM and dipped into ARMH3 (15-100 nM). All curves were fit with a partial, 1:1 binding model. The K_D_ value reported was generated using the average K_D_ given for each curve meeting inclusion criteria, error is reported as standard deviation (n = 5). **E)** Sum of the number of deuteron differences in ARMH3 upon binding to Arl5B(GTPγS) analyzed over the entire deuterium exchange time course. Each point represents the centre residue of an individual peptide. Peptides that met the significance criteria (defined as >0.4 Da, >5%, and *p* <0.01 in an unpaired two-tailed *t* test at any time point) are coloured red. Error is shown of the sum of standard deviations (SDs) across all five time points (n = 3). **F)** Selected deuterium exchange time courses that showed significant decreases in exchange upon complex formation. Error is shown as SD (n = 3). **G)** AlphaFold3 prediction of ARMH3 in complex with Arl5B and co-factors GTP and Mg_2+_ coloured by chain, important structural features of ARMH3 and Arl5B are annotated. **H)** Predicted aligned error (pae) plot of the AlphaFold3 prediction of the ARMH3-Arl5B(GTP) complex with co-factors GTP and Mg_2+_. **I)** AlphaFold3 model of the ARMH3-Arl5B complex coloured by significant decreases in deuterium exchange in ARMH3 upon binding to Arl5B(GTPγS). Boxes highlight significant peptides at the ARMH3-Arl5B interface and distal regions of ARMH3 showing allosteric changes.

To further understand the structural basis of the Arl5-ARMH3 interaction, we used hydrogen deuterium exchange mass spectrometry (HDX-MS), a powerful technique used to investigate protein conformational dynamics. It measures the exchange of backbone amide hydrogens with deuterium, with the rate of exchange being primarily dependent on secondary structure (Masson *et al*, 2017; James *et al*, 2022; Masson *et al*, 2019). HDX experiments were performed on ARMH3 alone and in complex with two-fold excess of Arl5B(GTPγS). Deuterium incorporation of pepsin-generated peptides was measured across five time points: 3s at 0°C, and 3s, 30s, 300s, 3000s at 18°C. The mass shift upon deuterium incorporation was analyzed via mass spectrometry, and for all HDX-MS experiments a change is considered significant when there was a >0.4 Da mass shift and >5% difference in percent deuterium incorporation and with *p* values less than 0.01 in a two-tailed unpaired *t* test at any time point. The full experimental details are present in the HDX stats table in the source data. Upon Arl5B(GTPγS) binding, we observed a significant decrease in deuterium incorporation at the N-terminus of ARMH3 (region spanning residues 11-63) and at the C-terminus (region spanning residues 674-689). Notably, we also saw a decrease in exchange spanning residues 623-642, towards the C-terminus but distinct from changes observed at the very C-terminus of ARMH3 (**Fig 1E+F**).

To interpret the HDX data on a molecular model, we modelled the interface of Arl5B with ARMH3 and GTP and Mg^2+^ using AlphaFold3 (Abramson *et al*, 2024, 3). This resulted in a high confidence prediction with low predicted aligned error between Arl5B and ARMH3 (**Fig 1G+H, Fig S1**). The HDX-MS differences matched well to the AlphaFold3 prediction of the ARMH3-Arl5B interaction. The N-terminus of ARMH3 engages with the switch I region of Arl5B (M38-S50). Interestingly, ARMH3 makes no contact with the switch II region (D66-T82), instead forming an interface with the C-terminus of Arl5B. ARMH3 itself is composed of N- to C-terminal armadillo repeat bundles, with a linker that runs across the armadillo repeat orienting the C-terminus next to the N-terminus, with α1 and α32 both showing decreased deuterium exchange at the Arl5B interface. The deuterium exchange difference in ARMH3 at residues 623-642 is distal to the Arl5 interface, indicating a possible allosteric change that propagates through the linker of ARMH3 upon Arl5 binding (**Fig 1I**).

To further validate the ARMH3-Arl5B(GTPγS) interface, we performed site-directed mutagenesis on key interface residues (**Fig 2A**). Examination of the predicted interface and evolutionary conservation of residues in both ARMH3 and Arl5B led us to generate mutations to interrupt complex formation (**Fig 2B**). Indeed, mutations along the N-terminus of ARMH3, including K24E, H93E, and K53E all abolished binding to Arl5B(GTPγS). Mutation of the residue in contact with H93 of ARMH3 in Arl5B (E163K) also led to a complete disruption of ARMH3 binding (**Fig 2C+D**).

**Figure 2:**
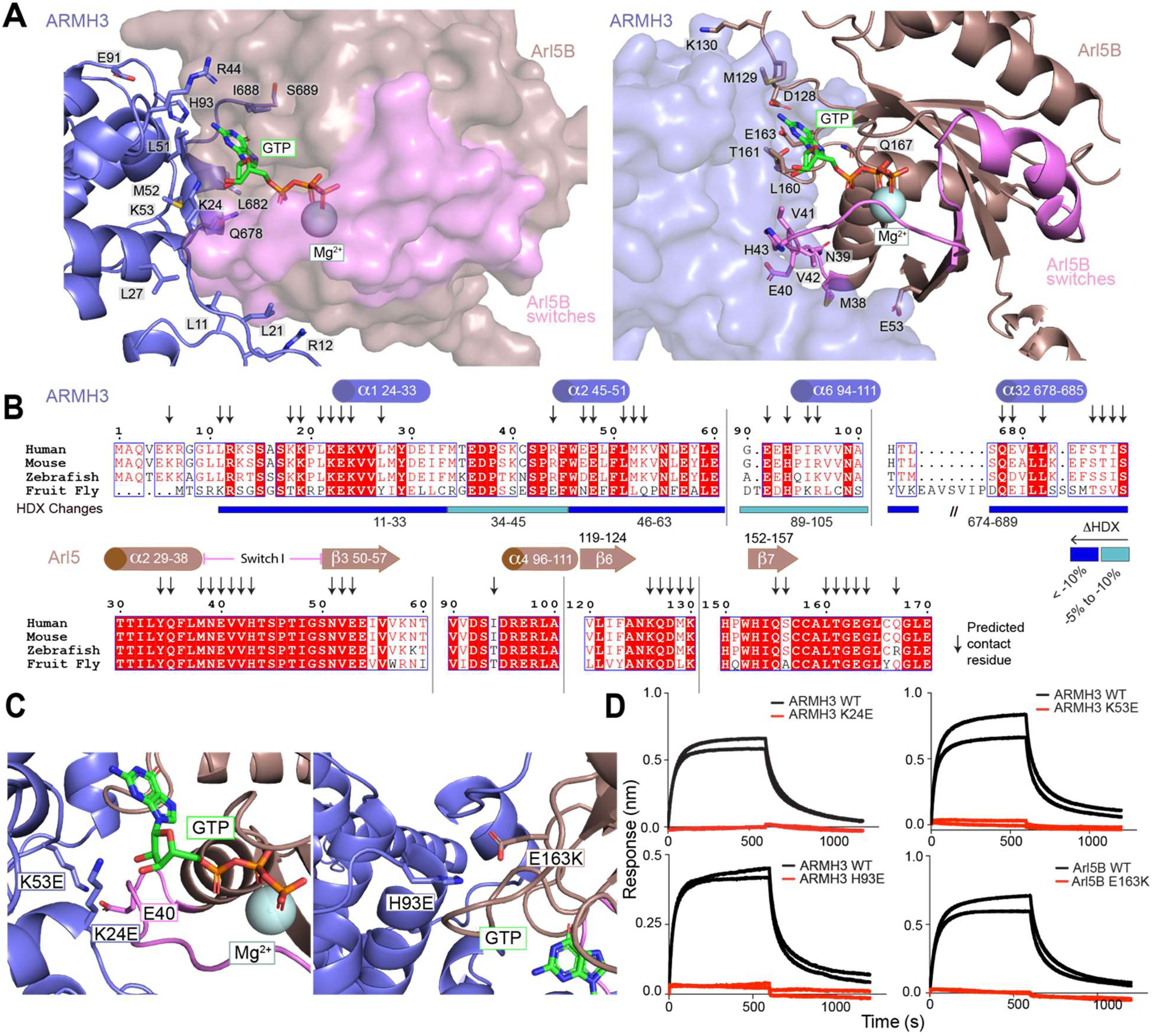
Molecular basis of Arl5B(GTP) binding to ARMH3. **A)** Zoom in on the predicted ARMH3-Arl5B(GTP) interface, showing select contact residues of ARMH3 and Arl5B labeled and shown as sticks. Switch regions of Arl5B are coloured pink. **B)** Multiple sequence alignments of ARMH3 and Arl5 from *H. sapiens* (human), *M. musculus* (mouse), *D. rerio* (zebrafish), and *D. melanogaster* (fruit fly). Secondary structures of ARMH3 and Arl5 are annotated above the alignment, while the HD exchange differences described in Fig. 1 are annotated below the ARMH3 alignment. Predicted contact residues with >5 Å^2^ of buried surface area are annotated using arrows. **C)** Zoomed in view of the ARMH3-Arl5B(GTP) interface with key residues and mutations labelled. **D)** Biolayer interferometry (BLI) association and dissociation curves of ARMH3 and Arl5B(GTPγS) mutants compared to wild-type. His-Arl5B(GTPγS) was loaded onto the anti-penta His tip at 25 nM and dipped into ARMH3 at 100 nM.

The allosteric changes observed in ARMH3 upon Arl5B binding are in a region that showed decreased exchange in ARMH3 upon complex formation with PI4KB (**Fig 3A**) (McPhail *et al*, 2020). Changes in H/D exchange reported in (McPhail *et al*, 2020) were re-analyzed using additional peptides from our analysis of Arl5 binding. The increased coverage of ARMH3 revealed previously uncharacterized changes upon PI4KB binding, including increased exchange from residues 183-190 and decreased exchange from residues 227-232 (**Fig 3B**). Re-analyzed data mapped onto the AlphaFold3 model of the ARMH3-PI4KB complex (**Fig 3C+D, Fig S2**) revealed a significant decrease in deuterium incorporation (>30 %D) in ARMH3 localised to the same region that exhibits decreased exchange upon Arl5B binding (residues 633-641). Interestingly, there was increased H/D exchange upon PI4KB binding at the Arl5B binding site on ARMH3, showing that Arl5 or PI4KB binding cause allosteric conformational changes to the other site. Changes in PI4KB mapped onto the ARMH3-PI4KB AlphaFold3 prediction reveal how the region with the largest decrease in deuterium incorporation in PI4KB (N-lobe linker of PI4KB, residues 488-515) interacts with ARMH3 (**Fig 3E**).

**Figure 3.**
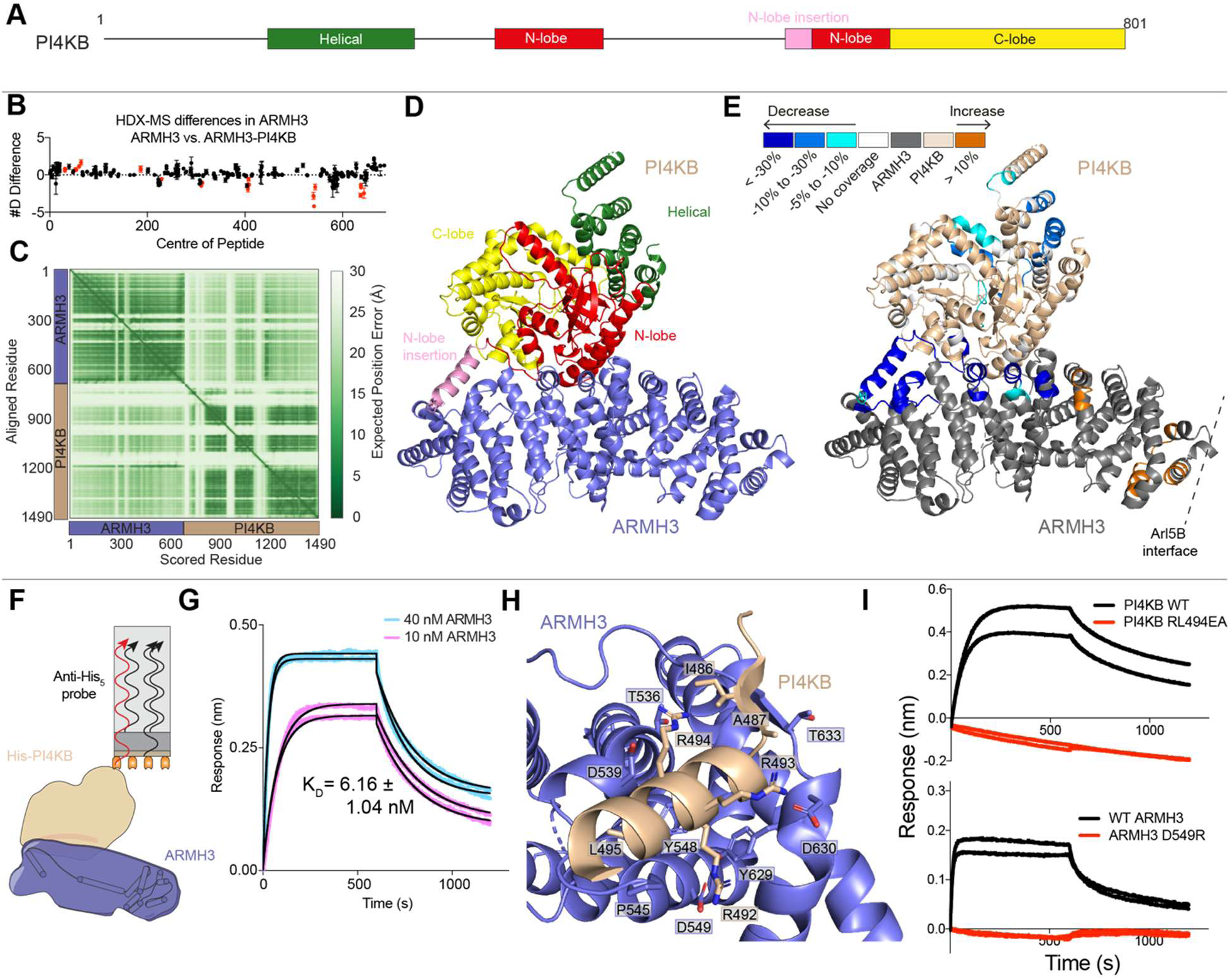
Molecular basis of the PI4KB-ARMH3 interaction. **A)** Domain schematic of the full-length PI4KB isoform 2 construct used in this paper. **B)** Sum of the number of deuteron differences in ARMH3 upon binding to PI4KB analyzed over the entire deuterium exchange time course, with additional ARMH3 peptides from ARMH3-Arl5B and ARMH3-GBF1 experiments included. Each point represents the centre residue of an individual peptide. Peptides that met the significance criteria (defined as >0.4 Da, >5%, and *p* <0.01 in an unpaired two-tailed *t* test at any time point) are coloured red. Error is shown of the sum of standard deviations (SDs) across all four time points (n = 3). **C)** Predicted aligned error (pae) plot of the AlphaFold3 prediction of the ARMH3-PI4KB complex. **D)** AlphaFold3 prediction of ARMH3 in complex with PI4KB, coloured by domain. **E)** AlphaFold3 prediction of ARMH3 in complex with PI4KB coloured by significant changes in deuterium exchange in both ARMH3 (grey) and PI4KB (wheat) upon complex formation. **F)** Schematic of the PI4KB-ARMH3 biolayer interferometry (BLI) assay, showing binding of immobilised His-PI4KB on the tip to full-length ARMH3 in solution. **G)** Dose response of PI4KB binding to ARMH3. His-PI4KB was loaded onto the anti-penta His tip at 50 nM and dipped into ARMH3 (10 nM, 40 nM). All curves were fit with a partial, 1:1 binding model. The K_D_ value reported was generated using the average K_D_ given for each curve meeting inclusion criteria, error is reported as standard deviation (n = 4). **H)** Zoom in on the predicted ARMH3-PI4KB N-lobe linker interface, showing select predicted contact residues of ARMH3 and PI4KB labeled and shown as sticks. **I)**BLI association and dissociation curves of ARMH3 and PI4KB mutants compared to wild-type. Either His-PI4KB or His-ARMH3 was loaded onto the anti-penta His tip at 50 nM and dipped into ARMH3 or PI4KB at 50 nM.

To further characterize the interaction between ARMH3 and PI4KB, we performed BLI experiments to determine affinity and validate the interface (**Fig 3F**). To measure the kinetics of the interaction, PI4KB was purified as described previously (McPhail *et al*, 2020), and BLI experiments were performed using immobilised His-tagged PI4KB (50 nM), which was dipped into ARMH3 at 10 nM and 40 nM. The dissociation constant for the ARMH3-PI4KB interaction was 6.16 ± 1.04 nM, calculated as the mean K_D_ across all curves meeting the inclusion criteria (**Fig 3G**). Average binding parameters (K_D_, K_on_, K_off_) are shown in Table S1, with data and binding parameters for each individual curve available in the source data. Site-directed mutagenesis was performed to validate key residues in the predicted PI4KB-ARMH3 interface (**Fig 3H**). A D549R mutation in ARMH3 that is located at a D549-R492 salt bridge between ARMH3 and PI4KB completely eliminated binding to PI4KB. This prediction was consistent with a previously identified PI4KB RL494EA mutant (McPhail *et al*, 2020), with this also preventing PI4KB-ARMH3 complex formation (**Fig 3I**).

To further investigate the different known ARMH3 binding partners at the Golgi, we next focused our attention on the Arf1 GEF GBF1. We were unable to produce full length GBF1 from *Sf9* insect cells, so we attempted recombinant expression of multiple truncations of GBF1, with the construct spanning residues 1-709 yielding the highest amount of homogenous protein as determined by size exclusion chromatography and SDS-PAGE (**Fig 4A**). While we were able to generate protein, yields were low, preventing our ability to set a full deuterium exchange time course or use BLI to measure binding. To investigate the dynamics of the ARMH3-GBF1 interaction, we set a limited HDX experiment using full-length ARMH3 and GBF1(1-709) with apo GBF1 and in complex with ARMH3 incubated with D_2_O buffer for 3 s at 0°C. We observed a significant decrease in deuterium incorporation in GBF1 when bound to ARMH3 in a single region spanning residues 377-385 (**Fig 4B**). To interpret these differences with a molecular model, we ran AlphaFold3 predictions of dimeric 1-709 of GBF1 bound to two ARMH3 molecules (**Fig 4C+D**). AlphaFold3 predicted a stable dimeric interface between the two GBF1 molecules with low predicted alignment error. For binding to ARMH3, there was only a single region of GBF1 with a high confidence prediction, with this being the region spanning residues 364-395 of GBF1. We repeated these AlphaFold3 searches with full length GBF1, with the region from 364-395 also showing low pae scores to ARMH3, and no other regions of ARMH3 predicted to bind GBF1 (**Fig S3**). As such, we have defined this disordered linker region located in the DCB-HUS domains region of GBF1 (residues 364-395) as the ARMH3 binding region (ABR). We mapped the HDX-MS differences of GBF1 1-709 binding to ARMH3, with the region with decreased exchange mapping on to the ABR (**Fig 4E**).

**Figure 4:**
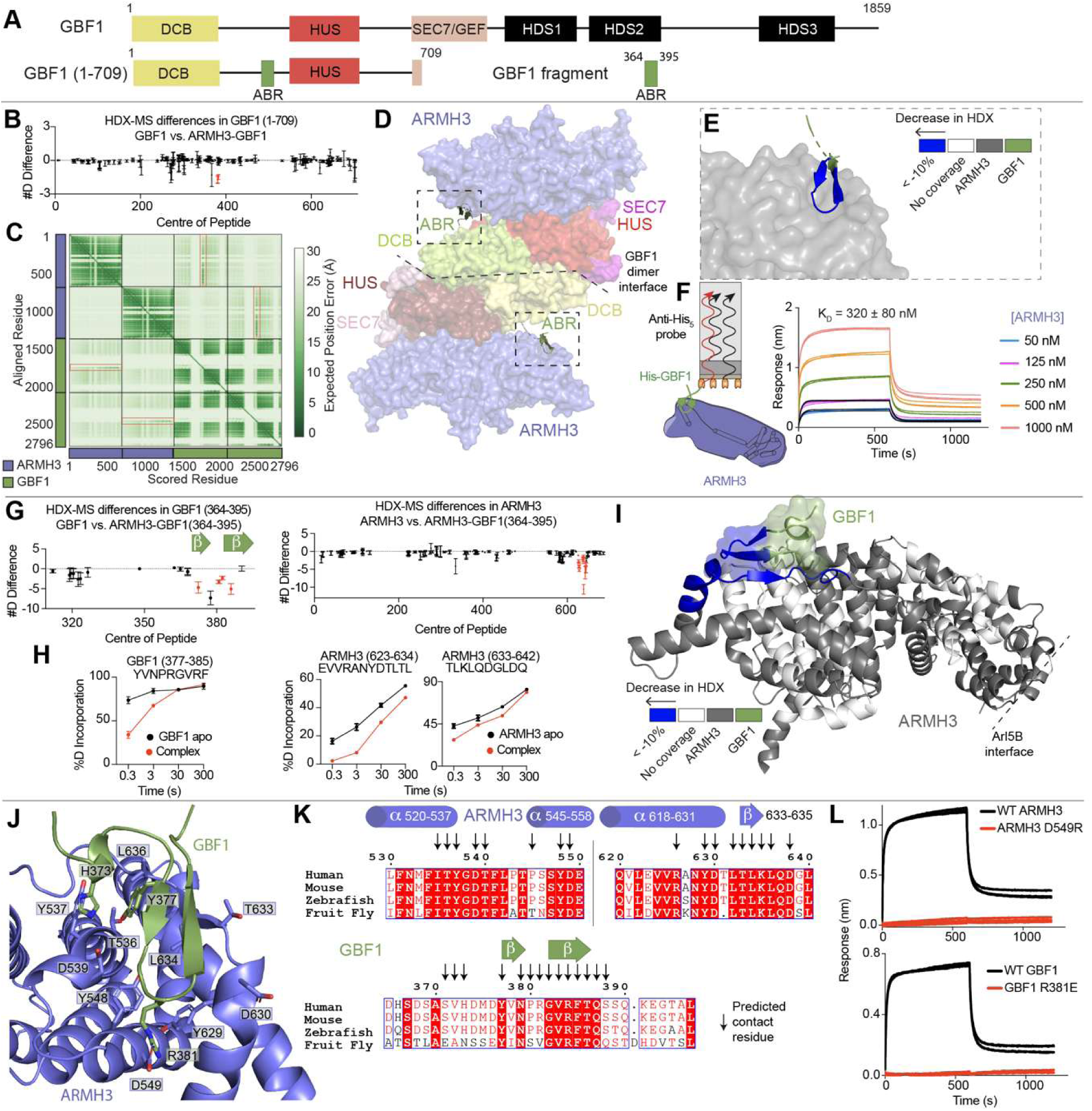
Molecular basis of GBF1 binding to ARMH3. **A)** Domain schematic of full-length GBF1 and the two constructs used in this paper, GBF1(1-709) and GBF1(364-395). **B)** Sum of the number of deuteron differences in GBF1(1-709) upon binding to ARMH3 analyzed over the 0.3s (3s at 0°C) deuterium exchange time course. Each point represents the centre residue of an individual peptide. Peptides that met the significance criteria (defined as >0.4 Da, >5%, and *p* <0.01 in an unpaired two-tailed *t* test) are coloured red. Error is shown of the sum of standard deviations across the 0.3s time point (n = 3). **C)** Predicted aligned error (pae) plot for the AlphaFold3 prediction of the ARMH3-GBF1(1-709) complex. The region corresponding to the predicted GBF1(364-395)-ARMH3 interface is boxed in red. **D)** AlphaFold3 prediction of ARMH3 in complex with GBF1(1-709) coloured by domain. The GBF1 ARMH3 binding region (ABR, residues 364-395) is shown as a cartoon. **E)** Zoom-in of the ABR-ARMH3 interface, coloured by significant changes in deuterium exchange in GBF1 upon ARMH3 binding. **F)** Schematic of the GBF1(364-395) biolayer interferometry (BLI) assay, showing binding of immobilised His-GBF1(364-395) binding to full-length ARMH3, and dose response curves of GBF1(364-395) binding to full-length ARMH3. His-GBF1(364-395) was loaded onto the anti-penta His tip at 100 nM and dipped into ARMH3 (50-1000 nM). All curves were fit with a partial, 1:1 binding model. The K_D_ value reported was generated using the average K_D_ given for each curve meeting inclusion criteria, error is reported as standard deviation (n = 4). **G)** Sum of the number of deuteron differences in GBF1(364-395) and ARMH3 upon binding to ARMH3 or GBF1(364-395) analyzed over the entire deuterium exchange time course. Each point represents the centre residue of an individual peptide. Peptides that met the significance criteria (defined as >0.4Da, >5%, and *p* <0.01 in an unpaired two-tailed *t* test) are coloured red. Error is shown of the sum of standard deviations across all four time points (n = 3). **H)** Selected deuterium exchange time courses that showed significant decreases in exchange upon complex formation. Error is shown as SD (n = 3). **I)** AlphaFold3 prediction of the ARMH3-GBF1(364-395) complex coloured by significant changes in deuterium exchange in both ARMH3 (grey) and GBF1(364-395) (smudge) upon complex formation. **J)** Zoom in on the predicted ARMH3-GBF1(364-395) interface, showing select predicted contact residues of ARMH3 and GBF1 labeled and shown as sticks. **K)** Multiple sequence alignments of ARMH3 and GBF1 from *H. sapiens* (human), *M. musculus* (mouse), *D. rerio* (zebrafish), and *D. melanogaster* (fruit fly). Secondary structures of ARMH3 and GBF1 are annotated above the alignment. Predicted contact residues with >5 Å^2^ of buried surface area are annotated using arrows. **L)** BLI association and dissociation curves of ARMH3 and GBF1(364-395) mutants compared to wild-type. His-GBF1(364-395) was loaded onto the anti-penta His tip at 100 nM and dipped into ARMH3 at 250 nM.

Given the difficulty of expressing and purifying GBF1(1-709), subsequent experiments on the ARMH3-GBF1 interaction were performed using the isolated ABR of GBF1, which expresses well in *E. coli* and is homogenous, as determined by size exclusion chromatography and SDS-PAGE. To characterize the interaction between GBF1(364-395) and ARMH3, we performed BLI experiments to measure affinity. Experiments were performed using immobilised His-tagged GBF1(364-395) (100 nM), which was dipped into ARMH3 at varying concentrations (50-1000 nM). From this data, we determined the dissociation constant of the ARMH3-GBF1(364-395) interaction to be 320 ± 80 nM, calculated as the mean K_D_ across all curves meeting the inclusion criteria (**Fig 4F**). Average binding parameters (K_D_, K_on_, K_off_) are shown in Table S1, raw data and binding parameters for each individual curve are available in the source data.

To supplement HDX data from ARMH3-GBF1(1-709) experiments, we performed HDX-MS on ARMH3-GBF1(364-395) with and without ARMH3 at four time points of exchange (3s at 0°C, 3s, 30s, and 300s at 18°C). We observed significant decreases in deuterium exchange within ARMH3 throughout the entire ABR (residues 377-385) (**Fig 4G+H**). Mapped changes on the AlphaFold3 prediction of the ARMH3-GBF1(364-395) complex (**Fig S4**) show decreased exchange localised to residues 623-642 within ARMH3 (**Fig 4I**). Notably, this is the same region of ARMH3 that shows decreased deuterium incorporation upon PI4KB binding and undergoes an allosteric change upon Arl5B(GTPγS) binding. An examination of the predicted interface and sequence alignment between ARMH3 and GBF1(364-395) revealed conserved residues that are likely required for binding (**Fig 4J+K**). There is a predicted salt bridge between D549 of ARMH3 and R381 in GBF1, with charge reversal mutations in either protein completely blocking complex formation (**Fig 4L**). The disruption of both GBF1(364-395)-ARMH3 and PI4KB-ARMH3 complexes upon mutation of ARMH3 D549R suggests that PI4KB and GBF1 share an interface on ARMH3.

Given the HDX-MS data showing that PI4KB and GBF1 share an interface on ARMH3, we investigated whether these interactions may be mutually exclusive. Overlaying the ARMH3-PI4KB and ARMH3-GBF1(364-395) AlphaFold3 predictions showed that they occupy the same binding pocket, with D549 in ARMH3 making a key interaction with either R381 in GBF1 or R492 in PI4KB (**Fig 5A**). We carried out a competition experiment with GBF1(364-395), PI4KB, and ARMH3, where we investigated whether the addition of PI4KB could abolish the ARMH3-GBF1(364-395) interaction (**Fig 5B**). Immobilised His-GBF1(364-395) (100 nM) was loaded, and dipped into wells containing ARMH3 (250 nM) and increasing concentrations of PI4KB (25-250 nM). PI4KB diminished the GBF1(364-395)-ARMH3 response in a dose-dependent manner (**Fig 5C**). The ABR of GBF1 contains a phosphosite at Y377 (**Fig 5D**), for which phosphomimetic mutants lead to dramatically decreased Golgi recruitment (Walton *et al*, 2023). We generated a phosphomimetic Y377E variant of the ABR to test the role of PTMs in regulating the GBF1-ARMH3 complex. GBF1(364-395) Y377E had greatly decreased binding to ARMH3 (**Fig 5E**). Examining the predicted ARMH3 binding sites for both GBF1 and PI4KB show a relatively negative pocket (**Fig 5F**), explaining why phosphorylation can alter complex formation.

**Figure 5.**
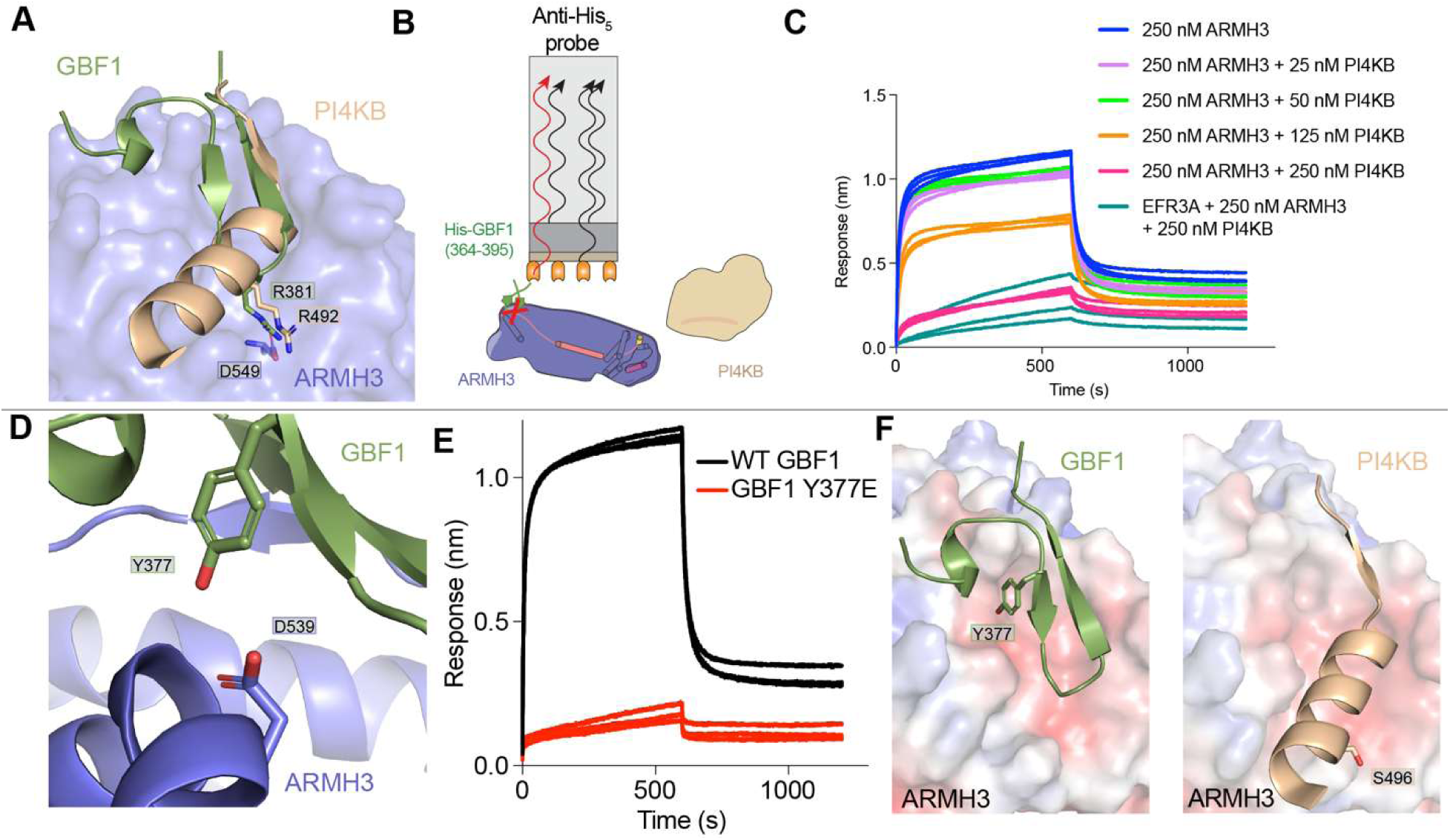
GBF1 and PI4KB phosphosites alter binding to ARMH3. **A)** Overlay of the AlphaFold3 predictions of PI4KB and GBF1 binding sites on ARMH3. ARMH3 is shown as a surface, GBF1 and PI4KB as cartoons, and important residues on GBF1 and PI4KB as sticks. **B)** Cartoon schematic of the GBF1(364-395)-ARMH3-PI4KB competition assay. His-tagged GBF1(364-395) was immobilised to the anti-His tip, dipping into 250 nM ARMH3 and increasing amounts of PI4KB (25-250 nM). **C)** Biolayer interferometry (BLI) traces of the GBF1(364-395)-ARMH3-PI4KB competition assay. **D)** Zoom-in of the Y377 phosphosite on GBF1, GBF1 Y377 and ARMH3 D539 are shown as sticks. **E)** BLI traces of WT GBF1 (100 nM) and phosphomimic mutant Y377E (100 nM) binding to ARMH3 (250 nM). **F)** GBF1(364-395)-ARMH3 and PI4KB N-lobe insertion (residues 483-500)-ARMH3 predicted interfaces with ARMH3 shown as an electrostatic surface. GBF1 phosphosite Y377 and PI4KB phosphosite S496 are shown as sticks.

## Discussion

ARMH3 has emerged as a critical scaffolding protein that plays important roles in the generation of both PI4P and active Arf1 at the Golgi (McPhail *et al*, 2020; Ishida *et al*, 2024; Chan *et al*, 2019). However, the molecular basis by which ARMH3 is recruited to the TGN, as well as how it recruits effectors, has remained unclear. Our biophysical and biochemical analysis of ARMH3 binding to Arl5, GBF1 and PI4KB has identified dynamic conformational changes, as well as the evolutionarily conserved binding interfaces to its targets. Overall, this provides fundamental insight into how ARMH3 recognizes its downstream effectors as well as how it can be targeted to the TGN by Arl5.

ARMH3 was originally identified as putative binding partner of PI4KB by affinity mass spectrometry (Greninger *et al*, 2012, 2013), with it found to be an essential gene (Blomen *et al*, 2015) that plays an important role in regulating PI4P levels at the Golgi/TGN (McPhail *et al*, 2020; Ishida *et al*, 2024; Mizuike *et al*, 2023). BioID experiments later revealed that ARMH3 interacts with the GEF GBF1, showing a potential impact on both Arf1 and PI4P at the Golgi (Chan *et al*, 2019). Recently, the GTPase Arl5 was identified as a potential regulator of PI4KB (Li *et al*, 2022), with this determined to be mediated by an interaction between ARMH3 and Arl5, with this playing an important role in regulating a distinct pool of PI4P at the TGN (Ishida *et al*, 2024). Our detailed biochemical and biophysical analysis has revealed the molecular basis for how Arl5 interacts with ARMH3. GTPγS-loaded Arl5 bound to ARMH3 with an affinity of ∼75 nM affinity, with the interface composed of both the N- and C-termini of ARMH3. AlphaFold3 (Abramson *et al*, 2024) modeling combined with HDX-MS analysis of the complex revealed potential interfacial residues, with complex disrupting mutations identified in both ARMH3 and Arl5. This interface appears to be strongly conserved through evolution, highlighting a potentially important role of this complex throughout multiple organisms. Intriguingly, Arl5 binding induced long-range allosteric conformational changes in ARMH3 at a region that had previously been suggested as the PI4KB binding site (McPhail *et al*, 2020), suggesting that TGN recruitment by Arl5 could prime ARMH3 for downstream effector engagement.

To understand how ARMH3 can carry out its roles once it is recruited by Arl5 required a molecular understanding of how it can interact with both GBF1 and PI4KB. Our HDX-MS and AlphaFold3 analysis of ARMH3 binding to PI4KB and GBF1 showed that they compete for a shared negative binding pocket in ARMH3. The ARMH3 interface in PI4KB is primarily centered around a linker region in the N-lobe of the kinase domain, although extensive additional conformational changes were also present in the kinase and helical domains of PI4KB that did not match well to the AlphaFold3 model. Further structural analysis will be required to understand how these possible allosteric conformational changes alter PI4KB regulation. In GBF1 there is a single disordered loop that binds to ARMH3, compared to the more extensive interface with PI4KB. Concurrently this likely explains why the interface with PI4KB has significantly higher affinity (∼6 nM) to ARMH3 compared to the weaker interface with GBF1 (∼300 nM). We identified a conserved residue in ARMH3 (D549), that forms critical salt bridges with R492 of PI4KB or R381 of GBF1. This suggests that ARMH3 is likely regulating distinct pools of GBF1 and PI4KB. This would be consistent with the idea that GBF1 is primarily localised at the Cis-Golgi (Manolea *et al*, 2008; Szul *et al*, 2007), while PI4KB and Arl5 are primarily at the TGN (Ishida *et al*, 2024; Mizuike *et al*, 2023). This suggests that potentially other upstream binding partners may interact with ARMH3 that control localisation to the Cis-Golgi, with BioID experiments with ARMH3 identifying multiple Cis-Golgi resident proteins that should be examined for direct binding to ARMH3 (Ishida *et al*, 2024).

Intriguingly, the regions of PI4KB and GBF1 binding interfaces to ARMH3 lie in regions that contain established phosphorylation sites. This suggests that phosphorylation could act as a switch to alter the recruitment of ARMH3 to either PI4KB or GBF1. We have previously shown that PKA phosphorylation of S496 (Isobe *et al*, 2017) in PI4KB leads to decreased affinity for ARMH3 (McPhail *et al*, 2020). Analysis of Golgi recruitment of phosphomimetic mutations of established phosphosites in GBF1 showed that Y377E led to the largest reduction in Golgi recruitment of all studied phosphosites (Walton *et al*, 2023). Our HDX-MS and AlphaFold3 modeling showed the Y377 residue makes a key interaction with ARMH3, with it pointing towards the acidic residue D539. The Y377E mutation completely abolished ARMH3 interaction, providing a molecular basis for why it leads to decreased Golgi recruitment of GBF1. One of the main unexplored aspects of this study is the potential set of kinases that could regulate this site. Recent advancements in determining kinase-substrate relationships (Yaron-Barir *et al*, 2024) allowed us to examine possible kinases that target Y377. The top five kinases were the pro-growth and immune kinases VEGFR1, ZAP70, JAK3, FAK, and FLT3. Most of these kinases are primarily plasma membrane localised, so further investigation will be required to understand how Y377 phosphorylation of GBF1 is controlled. Overall, this highlights how kinase-dependent PTMs located at this competitive binding interface could selectively route ARMH3 toward either PI4P synthesis or Arf1 activation, allowing dynamic redistribution of TGN signalling outputs without requiring changes in scaffold abundance.

The mechanistic framework here that defines how ARMH3 binds to members of the PI4P and Arf1 signalling cascades has direct implications for understanding how positive-strand RNA viruses generate replication organelles (van der Schaar *et al*, 2016; McPhail & Burke, 2023). Both GBF1 and PI4KB are essential host factors for multiple picornaviruses, where PI4P and GBF1 generated at viral replication organelles mediates lipid transport of cholesterol through regulation of lipid transfer proteins (van der Schaar *et al*, 2016; Ilnytska *et al*, 2013). Multiple picornaviruses utilise direct binding between their 3A proteins to GBF1, PI4KB or their adaptor proteins, including the PI4KB scaffolding protein ACBD3 (McPhail *et al*, 2017; Greninger *et al*, 2012; Lanke *et al*, 2009; Lyoo *et al*, 2019), to activate and recruit these enzymes to viral replication organelles. ARMH3 is required for viral replication in some enteroviruses, however, it is dispensable for others (McPhail *et al*, 2020; Blomen *et al*, 2015). This implies that some viruses may have evolved to directly manipulate ARMH3 for PI4KB activation, possibly as a mechanism to bypass ACBD3 mediated PI4KB activation. Further analysis of 3A proteins from viruses that require ARMH3 will be required to test this hypothesis.

Generation of PI4P and Arf1 at the Golgi is an essential process in Golgi maintenance and Golgi signalling. Our identification of how ARMH3 can play a critical scaffolding role in both PI4P or Arf1 signalling provides a key piece in the puzzle of understanding critical scaffolding roles at the Golgi. The identification of phosphoswitches in PI4KB and GBF1 at the ARMH3 interface reveal how ARMH3 interactions can be altered by kinase signalling. Our data highlight the molecular basis for complex formation of ARMH3 with Arl5, PI4KB, and GBF1, and how they have been conserved over evolution. Critically, ARMH3, Arl5, PI4KB, and GBF1 mutants developed in this study can be used in future *in vivo* experiments to study the role of ARMH3 at the Golgi and in positive-strand RNA virus replication.

### Experimental procedures

#### Plasmids

Plasmid containing GBF1(1-709), generated from GBF1 isoform 1 (UniProt Q92538-1, generously gifted by Dr. Chris Fromme, contains the QQ->Q variation and is one residue shorter than canonical Q92538-4) was cloned into a pLIB vector containing a 2x Strep and 10x His tag, followed by a TEV cleavage site (Gibson *et al*, 2009). Final pLIB constructs were transformed into DH10emBacY cells (Geneva Biotech) for blue-white screening; white colonies indicated successful bacmid generation containing the gene of interest. Original plasmids encoding Arl5A and Arl5B were sourced from DNASU, with accession numbers HsCD00002653 (Arl5A) and HsCD00042161 (Arl5B) (Seiler *et al*, 2014). Plasmids encoding Arl5A(15-179), Arl5B(15-179) and a truncated GBF1(364-395) were subcloned into a pOPT vector containing an N-terminal 2x Strep tag, followed by a 10x Histidine tag, followed by a tobacco etch virus (TEV) protease cleavage site (Gibson *et al*, 2009). Full length, wild-type 6x His-TEV ARMH3 and PI4KB WT and RL494EA was used as previously described (McPhail *et al*, 2020). Site-directed mutagenesis was performed on wild-type ARMH3, Arl5B, and GBF1(364-395) to generate mutant proteins for *E. coli* expression. The full list of all plasmids used in this study are shown in Table S2.

#### Protein expression

Plasmids containing the coding sequences for Arl5B(15-179), Arl5A(15-179), PI4KB RL494EA, GBF1(364-395), and both Arl5 and ARMH3 mutants were expressed in BL21 DE3 C41 *E. coli.* Cells were induced with 0.3 mM (Arl5) or 0.1 mM (ARMH3, PI4KB) isopropyl-β-D-thiogalactopyranoside (IPTG) and grown overnight at 16°C. Cells were then harvested and centrifuged at 1500xg, washed with PBS, and stored at −80°C. Bacmid harbouring wild-type ARMH3, PI4KB, and GBF1(1-709) were transfected into *Spodoptera frugiperda (Sf9)* cells and viral stock amplified for one generation to acquire a P2 generation final viral stock. Final viral stocks were added to 1-4 L of cells at a density of 2.0×10^6^ cells/mL in a 1:100 or 1:50 virus volume to cell volume ratio. Constructs were expressed for 55 to 68h before harvesting of the infected cells. Cell pellets were washed with PBS, flash-frozen in liquid nitrogen, and stored at −80°C.

#### Protein Purification – ARMH3

Cell pellets were lysed by sonication for 5 minutes in lysis buffer [20 mM tris (pH 8.0), 100 mM NaCl, 5% (v/v) glycerol, 20 mM imidazole, 2 mM β-mercaptoethanol (bME), and protease inhibitors (Millipore Protease Inhibitor Cocktail Set III, EDTA free)]. Triton X-100 was added to 0.1% (v/v), and the solution was centrifuged for 45 min at 20,000*g* at 1°C (Beckman Coulter J2-21, JA-20 rotor). The supernatant was then loaded onto a 5 mL HisTrap column (Cytiva) that had been equilibrated in nickel–nitrilotriacetic acid (Ni-NTA) A buffer [20 mM tris (pH 8.0), 100 mM NaCl, 20 mM imidazole (pH 8.0), 5% (v/v) glycerol, and 2 mM bME]. The column was washed with 4 column volumes (CV) of high-salt buffer [20 mM tris (pH 8.0), 1 M NaCl, 5% (v/v) glycerol, and 2 mM bME], 4 CV of Ni-NTA A buffer, and 4 CV of 6% Ni-NTA B buffer [20 mM tris (pH 8.0), 100 mM NaCl, 200 mM imidazole (pH 8.0), 5% (v/v) glycerol, and 2 mM bME] before being eluted with 4 CV of 100% Ni-NTA B. Protein was concentrated in a 30 kDa molecular weight cutoff (MWCO) concentrator (Millipore Sigma). The His tag was cleaved with TEV protease containing a stabilising lipoyl domain (Lip-TEV). TEV cleavage proceeded overnight at 4°C, following which the protein was loaded onto the Superdex 200 Increase 10/300 GL (Cytiva) pre-equilibrated in GFB [20 mM HEPES (pH 7.5), 150 mM NaCl, 10% (v/v) glycerol, and 0.5 mM TCEP]. Proteins eluted off gel filtration at volumes consistent with a monomer. Fractions from a single peak were collected and concentrated in 30 kDa MWCO concentrator (Millipore Sigma), flash-frozen in liquid nitrogen, and stored at −80°C until further use.

#### Protein Purification – Arl5

Cell pellets were lysed by sonication for 5 minutes in lysis buffer [20 mM tris (pH 8.0), 100 mM NaCl, 5% (v/v) glycerol, 20 mM imidazole, 2 mM β-mercaptoethanol (bME), and protease inhibitors (Millipore Protease Inhibitor Cocktail Set III, EDTA free)]. Triton X-100 was added to 0.1% (v/v), and the solution was centrifuged for 45 min at 20,000*g* at 1°C (Beckman Coulter J2-21, JA-20 rotor). The supernatant was then loaded onto a 5 mL HisTrap column (Cytiva) that had been equilibrated in nickel–nitrilotriacetic acid (Ni-NTA) A buffer [20 mM tris (pH 8.0), 100 mM NaCl, 20 mM imidazole (pH 8.0), 5% (v/v) glycerol, and 2 mM bME]. The column was washed with 4 column volumes (CV) of high-salt buffer [20 mM tris (pH 8.0), 1 M NaCl, 5% (v/v) glycerol, and 2 mM bME], 4 CV of Ni-NTA A buffer, and 4 CV of 6% Ni-NTA B buffer [20 mM tris (pH 8.0), 100 mM NaCl, 200 mM imidazole (pH 8.0), 5% (v/v) glycerol, and 2 mM bME] before being eluted with 4 CV of 100% Ni-NTA B. Protein was concentrated in a 10 kDa MWCO concentrator (Millipore Sigma) and buffer exchanged into GFB [20 mM HEPES (pH 7.5), 25 mM KCl, 5% (v/v) glycerol, 0.5 mM TCEP]. The protein was left overnight on ice at 4°C following which it was treated with 25 mM EDTA for 1 hour at 18°C before being centrifuged at 15,000*g* for 5 min. Protein was then buffer exchanged into phosphatase buffer [ 20 mM tris pH 8, 200 mM ammonium sulfate, 0.1 mM ZnCl_2_, 2 mM bME] before being treated with 2 units alkaline-agarose phosphatase (Millipore Sigma, P0762) per 1 mg protein. Beads were then separated from the protein using a 0.22 µm Millipore microcentrifuge filter, flowthrough was collected and spun for 1 minute at 15,000*g*. GDP or GTPγS was then added to 2- to 3-fold molar excess, and incubated for 1 hour at 18°C. Protein was then spun for 5 min at 15,000*g*. 30 mM MgCl_2_ was added to the protein solution, and was left to incubate for 20 min at 18°C. Protein was then loaded onto the Superdex 75 Increase 10/300 GL (Cytiva) pre-equilibrated in GFB + MgCl_2_ [20 mM HEPES (pH 7.5), 25 mM KCl, 5% (v/v) glycerol, 1 mM MgCl_2_, 0.5 mM TCEP]. Arl5B(GDP) eluted off gel filtration at volumes consistent with a dimer, whereas Arl5B(GTPγS) eluted off gel filtration at volumes consistent with a monomer. Fractions from a single peak were collected and concentrated in a 30 kDa MWCO concentrator (Millipore Sigma), flash-frozen in liquid nitrogen, and stored at −80°C until further use.

#### Protein Purification – PI4KB

Cell pellets were lysed by sonication for 3 minutes in lysis buffer [20 mM tris (pH 8.0), 100 mM NaCl, 5% (v/v) glycerol, 20 mM imidazole, 2 mM β-mercaptoethanol (bME), and protease inhibitors (Millipore Protease Inhibitor Cocktail Set III, EDTA free)]. Triton X-100 was added to 0.1% (v/v), and the solution was centrifuged for 45 min at 20,000*g* at 1°C (Beckman Coulter J2-21, JA-20 rotor). The supernatant was then loaded onto a 5 mL HisTrap column (Cytiva) that had been equilibrated in nickel–nitrilotriacetic acid (Ni-NTA) A buffer [20 mM tris (pH 8.0), 100 mM NaCl, 20 mM imidazole (pH 8.0), 5% (v/v) glycerol, and 2 mM bME]. The column was washed with 4 CV of high-salt buffer [20 mM tris (pH 8.0), 1 M NaCl, 5% (v/v) glycerol, and 2 mM bME], 2CV of Ni-NTA A buffer, then 1 CV of Ni-NTA A containing 2 mM adenosine 5′-triphosphate, 10 mM MgCl_2_, and 150 mM KCl. Column was then washed with 1 CV Ni-NTA A, followed by stepwise washes of 2 CV each with Ni-NTA B buffer in 10% increments; remaining volumes adjusted with Ni-NTA A. After the 70% Ni-NTA B fraction, the column was washed with 3 CV 100% Ni-NTA B to collect any remaining protein on the column. The 40%-100% Ni-NTA B fractions were collected and concentrated in a 50 kDa MWCO concentrator (Millipore Sigma) and buffer exchanged into GFB [20 mM HEPES (pH 7.5), 150 mM NaCl, 5% (v/v) glycerol, and 0.5 mM TCEP]. The His-tag was cleaved with Lip-TEV overnight at 4°C, following which the protein was loaded onto the Superdex 200 Increase 10/300 GL (Cytiva) pre-equilibrated in GFB. Proteins eluted off gel filtration at volumes consistent with a monomer. Fractions from a single peak were collected and concentrated in a 50 kDa MWCO concentrator (Millipore Sigma), flash-frozen in liquid nitrogen, and stored at −80°C until further use.

#### Protein Purification – GBF1(1-709)

Cell pellets were lysed by sonication for 3 minutes in lysis buffer [20 mM tris (pH 8.0), 100 mM NaCl, 5% (v/v) glycerol, 20 mM imidazole, 2 mM β-mercaptoethanol (bME), and protease inhibitors (Millipore Protease Inhibitor Cocktail Set III, EDTA free)]. Triton X-100 was added to 0.1% (v/v), and the solution was centrifuged for 45 min at 20,000g at 1°C (Beckman Coulter J2-21, JA-20 rotor). The supernatant was then loaded onto a 5 mL HisTrap column (Cytiva) that had been equilibrated in nickel–nitrilotriacetic acid (Ni-NTA) A buffer [20 mM tris (pH 8.0), 100 mM NaCl, 20 mM imidazole (pH 8.0), 5% (v/v) glycerol, and 2 mM bME]. The column was washed with 4 column volumes (CV) of high-salt buffer [20 mM tris (pH 8.0), 1 M NaCl, 5% (v/v) glycerol, and 2 mM bME], 4 CV of Ni-NTA A buffer, and 4 CV of 6% Ni-NTA B buffer [20 mM tris (pH 8.0), 100 mM NaCl, 200 mM imidazole (pH 8.0), 5% (v/v) glycerol, and 2 mM bME] before being eluted with 4 CV of 100% Ni-NTA B. The eluate was the loaded onto a 5mL StrepTrapHP column (Cytiva) and then washed with 3 CV of GFB [20 mM HEPES (pH 7.5), 150 mM NaCl, 10% (v/v) glycerol, 0.5 mM TCEP]. Protein was eluted with 3 CV of GFB containing 2.5 mM desthiobiotin and concentrated in a 50 kDA MWCO concentrator (Millipore Sigma). Concentrated protein was loaded onto the Superdex 200 Increase 10/300 GL (Cytiva) or the Superose 6 Increase 10/300 GL (Cytiva) pre-equilibrated in GFB. Proteins eluted off gel filtration at volumes consistent with a dimer. Fractions from a single peak were collected and concentrated in a 50 kDa MWCO concentrator (Millipore Sigma), flash frozen in liquid nitrogen, and stored at −80°C.

#### Protein Purification – GBF1(364-395)

Cell pellets were lysed by sonication for 5 minutes in lysis buffer [20 mM tris (pH 8.0), 100 mM NaCl, 5% (v/v) glycerol, 20 mM imidazole, 2 mM β-mercaptoethanol (bME), and protease inhibitors (Millipore Protease Inhibitor Cocktail Set III, EDTA free)]. Triton X-100 was added to 0.1% (v/v), and the solution was centrifuged for 45 min at 20,000g at 1°C (Beckman Coulter J2-21, JA-20 rotor). The supernatant was then loaded onto a 5 mL HisTrap column (Cytiva) that had been equilibrated in nickel–nitrilotriacetic acid (Ni-NTA) A buffer [20 mM tris (pH 8.0), 100 mM NaCl, 20 mM imidazole (pH 8.0), 5% (v/v) glycerol, and 2 mM bME]. The column was washed with 4 column volumes (CV) of high-salt buffer [20 mM tris (pH 8.0), 1 M NaCl, 5% (v/v) glycerol, and 2 mM bME], 4 CV of Ni-NTA A buffer, and 4 CV of 6% Ni-NTA B buffer [20 mM tris (pH 8.0), 100 mM NaCl, 200 mM imidazole (pH 8.0), 5% (v/v) glycerol, and 2 mM bME] before being eluted with 4 CV of 100% Ni-NTA B. The eluate was the loaded onto a 5mL StrepTrapHP column (Cytiva) and then washed with 3 CV of GFB [20 mM HEPES (pH 7.5), 150 mM NaCl, 5% (v/v) glycerol, 0.5 mM TCEP]. Protein was eluted with 3 CV of GFB containing 2.5 mM desthiobiotin and concentrated in a 3 kDA MWCO concentrator (Millipore Sigma). Concentrated protein was loaded onto the Superdex 75 Increase 10/300 GL (Cytiva) pre-equilibrated in GFB. Proteins eluted off gel filtration at volumes consistent with a monomer. Fractions from a single peak were collected and concentrated in a 3 kDa MWCO concentrator (Millipore Sigma), flash frozen in liquid nitrogen, and stored at −80°C.

#### AlphaFold3 Predictions

We used the protein prediction software AlphaFold3 (Abramson *et al*, 2024) to generate predictions of the ARMH3-Arl5B, ARMH3-GBF1, and ARMH3-PI4KB interfaces. Specifically, we used the AlphaFold server (https://alphafoldserver.com) and input the sequences for full-length human ARMH3, GBF1, PI4KB, and Arl5B. For the ARMH3-Arl5B prediction, cofactors GTP and Mg^2+^ were added. As GBF1 is an obligate dimer, two copies of ARMH3 and GBF1 were inputted into the server to generate a physiologically accurate prediction of the GBF1-ARMH3 interface. Predictions of the different GBF1 constructs bound to ARMH3 were also generated, i.e. 2xARMH3-2xGBF1(1-709) and ARMH3-GBF1(364-395). Given the variability of AlphaFold3 predictions, five individual searches were performed for each complex, using seed numbers 1 through 5. To evaluate the confidence of individual subunit assembly predictions, we analyzed the predicted alignment error (pae), predicted template modeling (pTM) score and the interface predicted template modeling (ipTM) score. We also analyzed the chain_pair_iptm and chain_pair_pae_min values. The chain_pair_iptm scores are useful in evaluating the confidence of predicted protein-protein interfaces, while the chain_pair_pae_min value correlates with whether two chains interact with each other. The prediction with the best scores were used to model the interfaces in this manuscript. chain_pair_iptm and chain_pair_pae_min scores for each search are available in supplementary data.

#### Sequence Alignments

Protein sequences from the UniProt data base were aligned using Clustal Omega Multiple Sequence Alignment and the aligned sequences were subsequently analyzed by ESPript 3.0 to visualize conserved regions. UniProt ARMH3 entries used: *H. sapiens* (Q5T2E6), *M musculus* (Q6PD19), *D. rerio* (Q6PGW3), *D. melanogaster* (Q7KSU3). UniProt Arl5 entries used: *H. sapiens* (Q96KC2), *M. musculus* (Q9D4P0), *D. rerio* (Q7SZE7), *D. melanogaster* (Q9VSG8). UniProt GBF1 entries used: *H. sapiens* (Q92538-1), *M. musculus* (Q6DFZ1), *D. rerio* (A0A8M3AYV9), *D. melanogaster* (A1Z8W8). The AlphaFold3 predicted protein interfaces were examined using the PDBePISA (Proteins, Interfaces, Structures and Assemblies) server (Krissinel & Henrick, 2007).

#### Biolayer Interferometry

The BLI measurements were conducted using a ForteBio (Sartorius) K2 Octet or GatorBio Prime Core BLI System using fiber optic biosensors. Anti-penta-His biosensors were loaded using either purified Arl5A, Arl5B, PI4KB, ARMH3, or GBF1(364-395), which all had a either a 6x or 10xHis tag on the N-terminus, used during protein purification. The biosensor tips were preincubated in the BLI buffer [ARMH3-Arl5, ARMH3-GBF1(364-395): 20 mM HEPES (pH 7.5), 25 mM KCl, 1 mM MgCl_2_, 0.01%, bovine serum albumin, and 0.002% Tween-20; ARMH3-PI4KB: 20 mM HEPES (pH 7.5), 150 mM NaCl, 0.01%, bovine serum albumin, and 0.002% Tween-20] for 10 min before experiments began. The sequence of steps in each assay was regeneration, custom, loading, baseline, association, and dissociation. For some experiments, a loading threshold of 0.5 nm was used to prevent nonspecific loading of tagged protein. Every experiment was done at 25°C with shaking at 1000 rpm. Technical replicates were performed using the same fiber tip and repeating the steps outlined previously. Regeneration was performed by exposing the tips to regeneration buffer (glycine pH 1.5 or GatorBio Regen Buffer) for 5s and BLI buffer for 5s and repeating the exposure for six cycles. BLI buffer was used for the custom, baseline, and dissociation steps; these steps were performed in the same well for a given sample. To determine binding kinetics, His-Arl5B(GTPγS) was diluted in BLI buffer to 25 nM, His-PI4KB to 50 nM, and His-GBF1(364-395) to 100 nM and were loaded onto the anti-penta-His biosensor tips. ARMH3 was also diluted in BLI buffer from 15 nM to 100 nM for Arl5B-ARMH3, 10 nM and 40 nM for PI4KB-ARMH3, and 50 nM to 1000 nM for GBF1-ARMH3 and added to the appropriate association wells. Data analysis was performed using the Octet Data Analysis HT software (Sartorius) or the GatorOne software (GatorBio). The data were fit using a 1:1 binding model using the “partial (each step separately)” setting. We only fit BLI curves that met the following criteria: a X^2^ value of less than 1.5, with a response >15% of the maximum fittable response. The kinetic binding constants (k_on_, k_off_, K_D_) were estimated as the mean of values from all such curves (ARMH3-Arl5B: n = 5, ARMH3-PI4KB: n = 4, ARMH3-GBF1(364-395): n = 4), with error reported as the standard deviation. The average kinetic binding data is reported in table S1, with all data for each curve shown in the source data.

For mutant binding experiments, Arl5B(GTPγS), PI4KB, and GBF1(364-395) wild-type or mutant was diluted in BLI buffer to 25 nM, 50 nM, and 100 nM, respectively. ARMH3 was diluted in BLI buffer to 100 nM (Arl5B-ARMH3), 50 nM (PI4KB-ARMH3) and 250 nM (GBF1(364-395)-ARMH3). These BLI experiments were performed using either two or three technical replicates.

For the competition experiment, GBF1(364-395) was diluted in BLI buffer to 100 nM and loaded onto the anti-His tips. ARMH3 was diluted to 250 nM, and PI4KB to 25 - 250 nM. PI4KB was added to the association wells in increasing concentrations (0, 25, 50, 125, and 250 nM). This BLI experiment was performed using three technical replicates.

#### HDX-MS sample preparation

HDX reactions comparing apo ARMH3 to ARMH3 incubated with Arl5B(GTPγS) (15-179) were carried out in a 50 µl reaction volume containing 15 pmol of ARMH3 and 30 pmol of Arl5B. The exchange reactions were initiated by the addition of 27.5 µL of D_2_O buffer (20 mM HEPES pH 7.5, 25 mM KCl, 1 mM TCEP, 1 mM MgCl_2_, 94.96% D_2_O (V/V)) to 22.5 µL of protein (final D_2_O concentration of 52.2%). Reactions proceeded for 3s on ice at 0°C (0.3s), as well as 3s, 30s, 300s, and 3000s at 18°C before being quenched with ice cold acidic quench buffer, resulting in a final concentration of 0.6M guanidine HCl and 0.9% formic acid post quench.

HDX reactions comparing apo ARMH3 to ARMH3 incubated with GBF1(1-709) were carried out in a 10.48 μL reaction volume; each experiment contained 20 pmol of both ARMH3 and GBF1(1-709). The exchange reactions were initiated by the addition of 5.83 μL D2O buffer (20 mM HEPES pH 7.5, 150 mM KCl, 0.5mM TCEP, 85.4% D2O (V/V)) to 4.65 μL of protein (final D2O concentration of 47.53%). Reactions proceeded for 3s at 0°C before being quenched with ice cold acidic quench buffer, resulting in a final concentration of 0.6M guanidine HCl and 0.9% formic acid post quench.

HDX reactions comparing apo ARMH3 to ARMH3 incubated with GBF1(364-395) were carried out in a 5 μL reaction volume. Each experiment contained 20 pmol of both ARMH3 and GBF1. The exchange reactions were initiated by the addition of 3 μL D2O buffer (20 mM HEPES pH 7.5, 150 mM NaCl, 0.5mM TCEP, 95.2% D2O (V/V)) to 2 μL of protein (final D2O concentration of 57.1%). Reactions proceeded for 3s at 0°C and 3s, 30s, and 300s at 18°C before being quenched with ice cold acidic quench buffer, resulting in a final concentration of 0.6M guanidine HCl and 0.9% formic acid post quench.

All conditions and time points were created and run in independent triplicate. Samples were flash frozen immediately after quenching and stored at −80°C until injected onto the ultra-performance liquid chromatography (UPLC) system for proteolytic cleavage, peptide separation, and injection onto a Orbitrap or QTOF for mass analysis, described below.

#### Protein Digestion and MS/MS Data Collection

Protein samples were rapidly thawed and injected onto an integrated fluidics system containing a HDx-3 PAL liquid handling robot and climate-controlled (2°C) chromatography system (LEAP Technologies), a Waters Acquity UPLC I-Class Series System, as well as an Orbitrap Exploris 120 Mass Spectrometer (ThermoFisher) (ARMH3-ARL5) or an Impact II QTOF Mass spectrometer (Bruker) (ARMH3-GBF1). The full details of the automated LC system were previously described in (Stariha *et al*, 2021). The samples were run over an immobilised pepsin column (Affipro; AP-PC-001) at 200 µL/min for 4 minutes at 2°C. The resulting peptides were collected and desalted on a C18 trap column (Acquity UPLC BEH C18 1.7 µm column (2.1 × 5 mm); Waters 186004629). For the Orbitrap Exploris, the trap was subsequently eluted in line with an ACQUITY 300Å, 1.7 μm particle, 100 × 2.1 mm BEH C18 UPLC column (Waters), using a gradient of 3-10% B (Buffer A 0.1% formic acid; Buffer B 100% acetonitrile) over 1.0 minutes, followed by a gradient of 10-25% B over 3.0 minutes, followed by a gradient of 25-35% B over 3 minutes, finally after 30 seconds at 35% B a gradient of 35-80% B over 1 minute was used. For the Impact II QTOF, the trap was eluted in line with an ACQUITY 300Å, 1.7 μm particle, 100 × 2.1 mm BEH C18 UPLC column (Waters), using a gradient of 3-10% B (Buffer A 0.1% formic acid; Buffer B 100% acetonitrile) over 1.5 minutes, followed by a gradient of 10-25% B over 4.5 minutes, followed by a gradient of 25-35% B over 5 minutes, finally after 1 minute at 35% B a gradient of 35-80% B over 1 minute was used. Mass spectrometry experiments acquired over a mass range from 150 to 2200 m/z using an electrospray ionization source operated at a temperature of 200°C and a spray voltage of 4.5 kV. Same LC system, gradient and columns were used for all samples.

#### Peptide identification

Peptides were identified from the non-deuterated samples of ARMH3 using data-dependent acquisition following tandem MS/MS experiments (0.5s precursor scan from 300-1500m/z, 4 data dependent scans from 300-1500m/z, dynamic exclusion set to n=1 with 10s exclusion time, 6s expected LC peak width). Peptides were identified from the non-deuterated samples ARMH3 and either GBF1(1-709) or GBF1(364-395) using data-dependent acquisition following tandem MS/MS experiments (0.5 s precursor scan from 150-2000 m/z; twelve 0.25 s fragment scans from 150-2000 m/z). MS/MS datasets were analyzed using FragPipe v18.0 (ARMH3-Arl5B) or FragPipe v23.1 (ARMH3-GBF1) and peptide identification was carried out by using a false discovery-based approach using a database of purified proteins and known contaminants (da Veiga Leprevost *et al*, 2020; Dobbs *et al*, 2020). MSFragger was used, and the precursor mass tolerance error was set from −20 to 20ppm. The fragment mass tolerance was set at 20ppm. Protein digestion was set as nonspecific, searching between lengths of 4 and 50 aa, with a mass range of 400 to 5000 Da (Kong *et al*, 2017).

#### Mass Analysis of Peptide Centroids and Measurement of Deuterium Incorporation

HD-Examiner Software (Sierra Analytics) was used to automatically calculate the level of deuterium incorporation into each peptide. All peptides were manually inspected for correct charge state, correct retention time, appropriate selection of isotopic distribution, etc. Deuteration levels were calculated using the centroid of the experimental isotope clusters. Results are presented as relative levels of deuterium incorporation and the only control for back exchange was the level of deuterium present in the buffer. Differences in exchange in a peptide were considered significant if they met all three of the following criteria: >5% change in exchange, ≥0.4 Da difference in exchange, and a *p* value <0.01 using a two tailed student *t*-test. Samples were only compared within a single experiment and were never compared to experiments completed at a different time with a different final D_2_O level. The data analysis statistics for all HDX-MS experiments can be found in the source data according to published guidelines (Masson *et al*, 2019). The mass spectrometry proteomics data have been deposited to the ProteomeXchange Consortium via the PRIDE partner repository (Perez-Riverol *et al*, 2021) with the dataset identifier (PXD076823).

#### Re-analysis of Peptide Centroids and Deuterium Incorporation in the PI4KB-ARMH3 dataset

Peptides identified as described above in non-deuterated samples of ARMH3 in the ARMH3-Arl5B and ARMH3-GBF1 experiments, as well as those generated in (McPhail *et al*, 2020), were compiled and used to find previously unidentifiable ARMH3 peptides in the PI4KB-ARMH3 dataset. HD-examiner software (Sierra Analytics) was used to automatically calculate the level of deuterium incorporation into each ARMH3 peptide. All peptides were manually inspected for correct charge state, correct retention time, appropriate selection of isotopic distribution, etc. Deuteration levels were calculated using the centroid of the experimental isotope clusters. Results are presented as relative levels of deuterium incorporation and the only control for back exchange was the level of deuterium present in the buffer (87%). Differences in exchange in a peptide were considered significant if they met all three of the following criteria: >5% change in exchange, ≥0.4 Da difference in exchange, and a p value <0.01 using a two tailed student t-test. Samples were only compared within the experiment and were never compared to experiments completed at a different time with a different final D_2_O level. The data analysis statistics for this re-analysis can be found in the source data according to published guidelines (Masson *et al*, 2019). The mass spectrometry proteomics data have been deposited to the ProteomeXchange Consortium via the PRIDE partner repository (Perez-Riverol *et al*, 2021) with the dataset identifier (PXD076708).

## Data availability

The MS proteomics data have been deposited to the ProteomeXchange Consortium via the PRIDE partner repository with the dataset identifier PXD076708 and PXD076823 (Perez-Riverol *et al*, 2021). All raw data in all figures are available in the source data excel file. All data needed to evaluate the conclusions in the paper are present in the paper and/or the supporting information.

## Author Contributions

MKS contributed to conceptualization, data curation, formal analysis, investigation, methodology, validation, visualization, writing – original draft, and writing – reviewing & editing. GCTK contributed to investigation, methodology, visualization, and writing – reviewing & editing. EEW contributed to data curation, formal analysis, methodology, resources, software, validation, visualization, and writing – reviewing & editing. SS contributed to data curation, formal analysis, methodology, validation, visualization, and writing – reviewing & editing. HGN contributed to data curation, formal analysis, validation, visualization, and writing – reviewing & editing. JEB contributed to conceptualization, funding acquisition, project administration, software, supervision, validation, visualization, writing – original draft, and writing – reviewing & editing.

## Supporting information

This article contains supporting information.

## Funding and additional information

J.E.B. is supported by a Natural Sciences and Engineering Research Council of Canada Discovery grant NSERC-2020-04241.

## Acknowledgements

The plasmid encoding GBF1 was a kind gift from the laboratory of Dr Chris Fromme. The plasmids encoding Arl5A and Arl5B were sourced from the DNASU depository run by Arizona State University.

## Supplementary Figures and Tables

**Figure S1:**
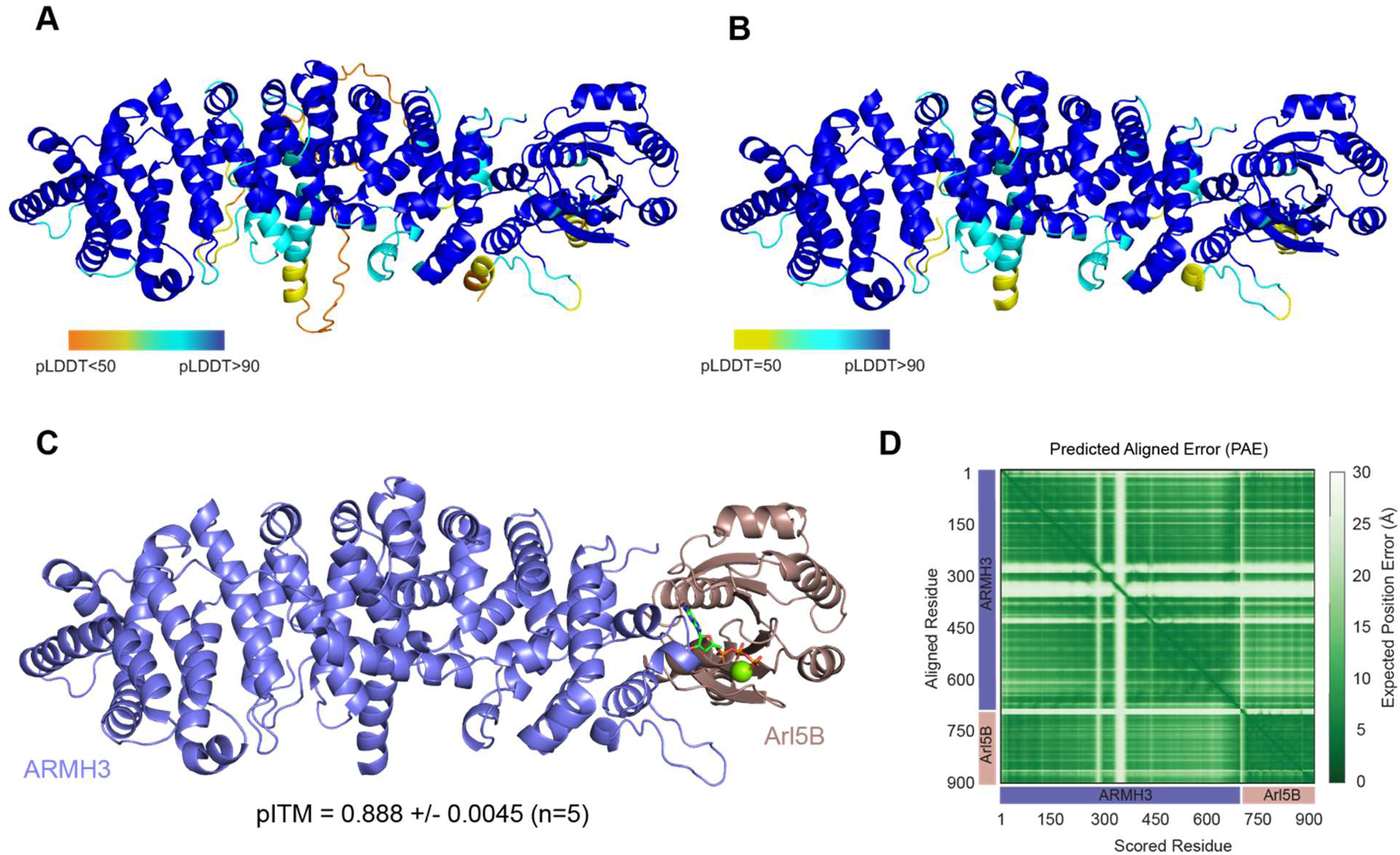
AlphaFold3 modeling of the ARMH3-Arl5B complex. **A)** Best model (seed 2, n=5) of the Arl5B-ARMH3 AlphaFold3 prediction with cofactors GTP and Mg^2+^ coloured by pLDDT. iPTM and chain_pair_pae_min information for individual seeds in source data. **B)** The same model coloured by pLDDT, with regions of low confidence removed. **C)** Model shown in panel B, coloured by chain. iPTM score represents the average across all five seeds, with error shown as standard deviation. **D)** Predicted aligned error (pae) plot for seed 2 of Arl5B in complex with ARMH3 and cofactors GTP and Mg^2+^.

**Figure S2:**
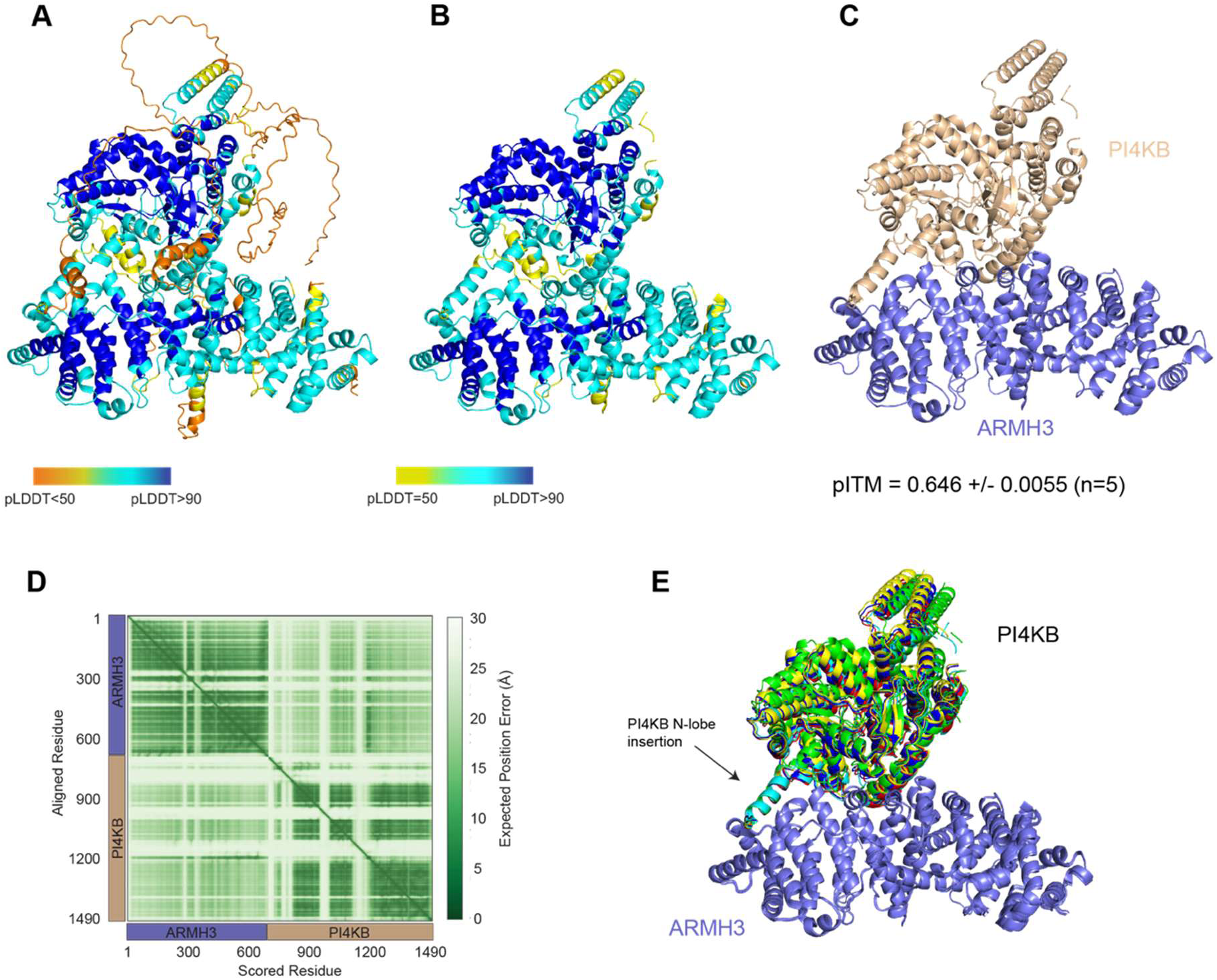
AlphaFold3 modeling of the PI4KB-ARMH3 complex. **A)** Best model (seed 5, n=5) of the ARMH3-PI4KB AlphaFold3 prediction coloured by pLDDT. iPTM and chain_pair_pae_min information for individual seeds in source data. **B)** The same model coloured by pLDDT, with regions of low confidence removed. **C)** Model shown in panel B, coloured by chain. iPTM score represents the average across all five seeds, with error shown as standard deviation. **D)** Predicted aligned error (pae) plot for seed 5 of PI4KB in complex with ARMH3. **E)** Alignment of all five seeds onto ARMH3; PI4KB N-lobe insertion indicated with arrow.

**Figure S3:**
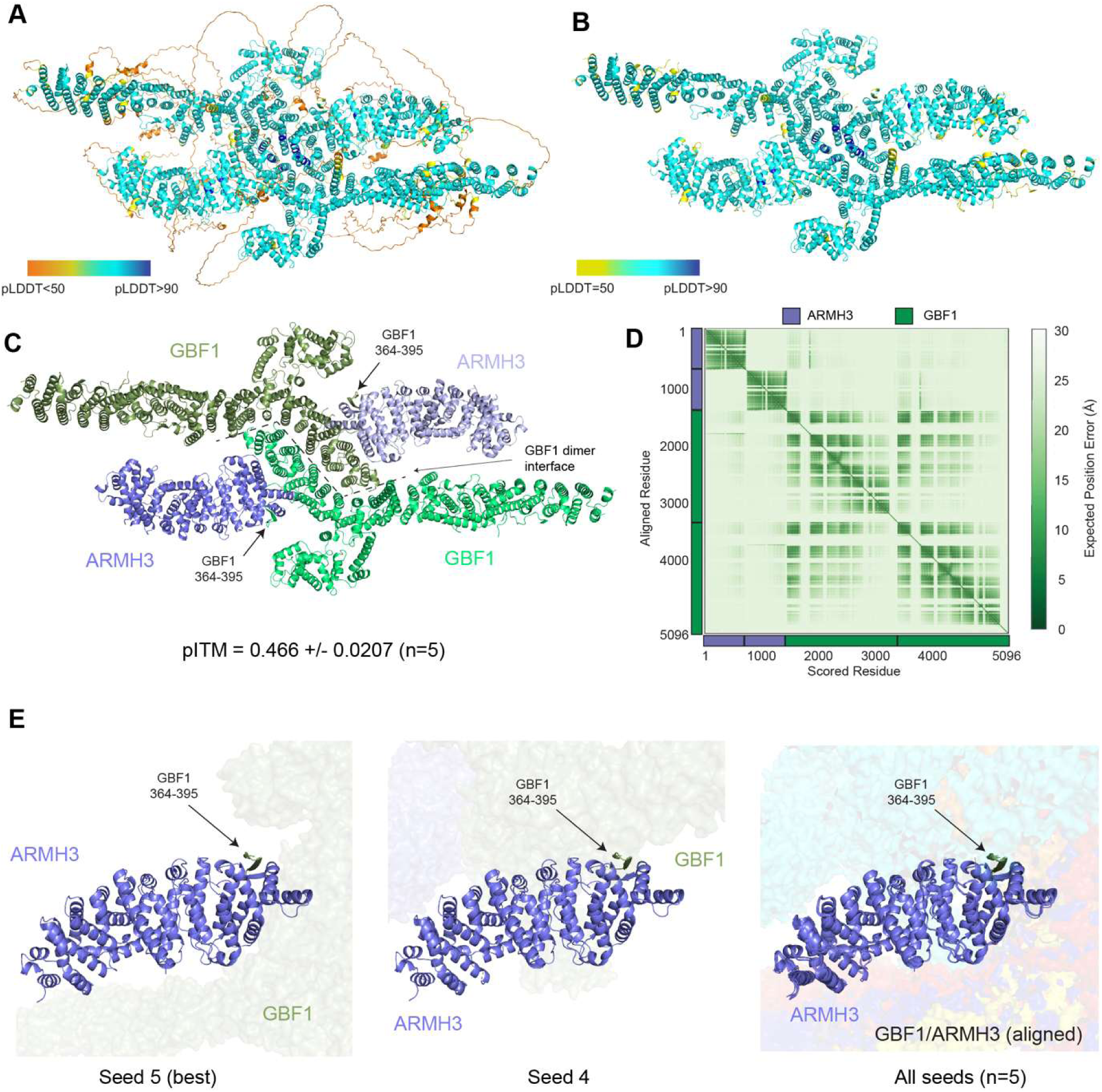
AlphaFold3 modeling of the ARMH3-GBF1 complex. **A)** Best model (seed 5, n=5) of the 2xARMH3-2xGBF1 AlphaFold3 prediction coloured by pLDDT. iPTM and chain_pair_pae_min information for individual seeds in source data. **B)** The same model coloured by pLDDT, with regions of low confidence removed. **C)** Model shown in panel B, coloured by chain. Predicted GBF1 dimer interface is indicated by the dashed line, arrows indicate GBF1 residues 364-395. iPTM score represents the average across all five seeds, with error shown as standard deviation. **D)** Predicted aligned error (pae) plot for seed 5 of 2xGBF1 with 2xARMH3. **E)** Zoom-ins showing predictions of ARMH3 in complex with full-length GBF1, with one ARMH3 and GBF1 residues 364-395 shown as cartoons and the remainder of the model shown as a surface. Left and middle panels show select predictions of the GBF1-ARMH3 interface, while the right panel shows all five predictions aligned onto ARMH3. GBF1 residues are indicated with an arrow.

**Figure S4:**
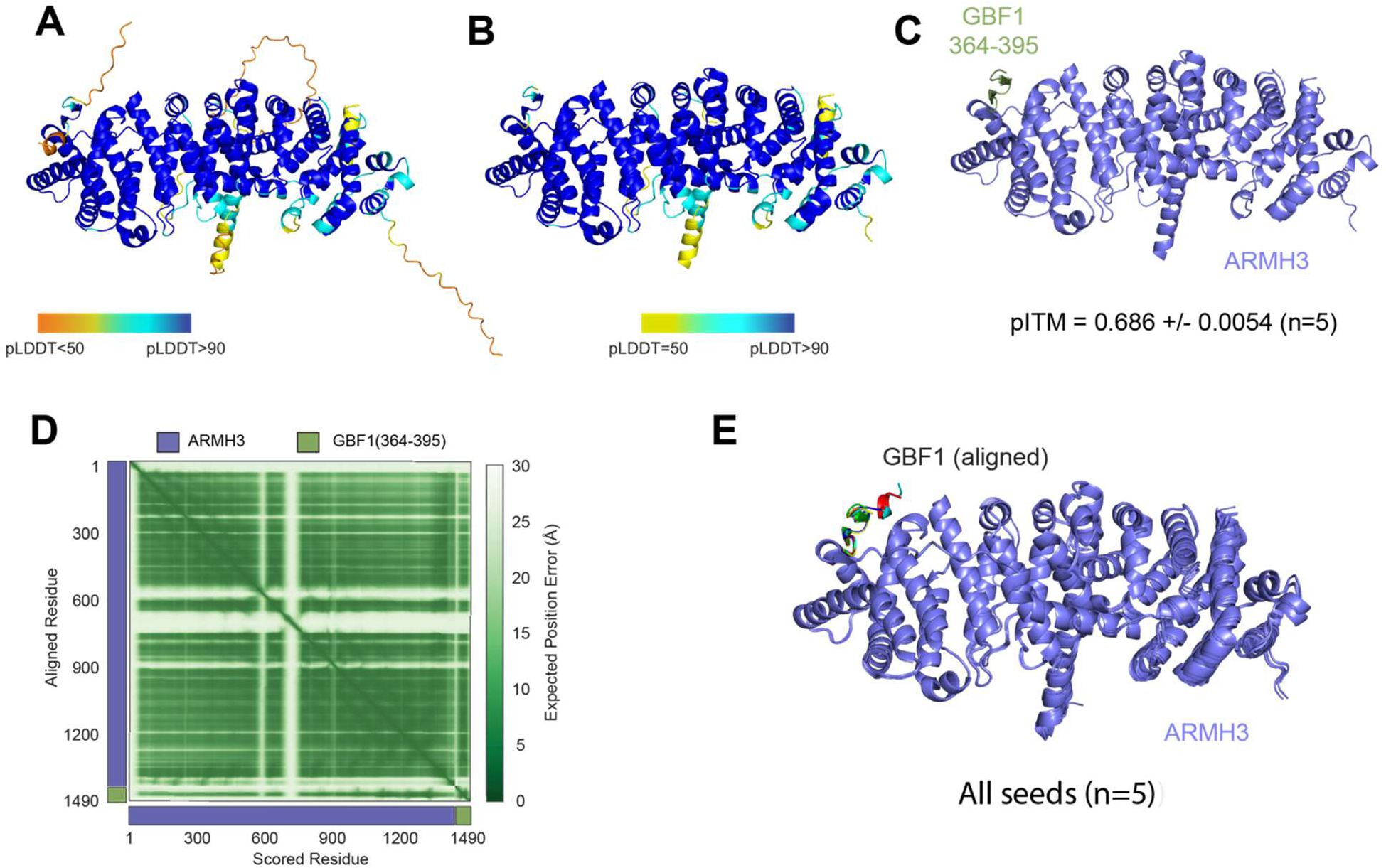
AlphaFold3 modeling of the ARMH3-GBF1(364-395) complex. **A)** Best model (seed 4, n=5) of the ARMH3-GBF1(364-395) AlphaFold3 prediction coloured by pLDDT. iPTM and chain_pair_pae_min information for individual seeds in source data. **B)** The same model coloured by pLDDT, with regions of low confidence removed. **C)** Model shown in panel B, coloured by chain. iPTM score represents the average across all five seeds, with error shown as standard deviation. **D)** Predicted aligned error (pae) plot for seed 4 of GBF1(364-395) in complex with ARMH3. **E)** Alignment of all five seeds onto ARMH3.

---

**Table S1.**
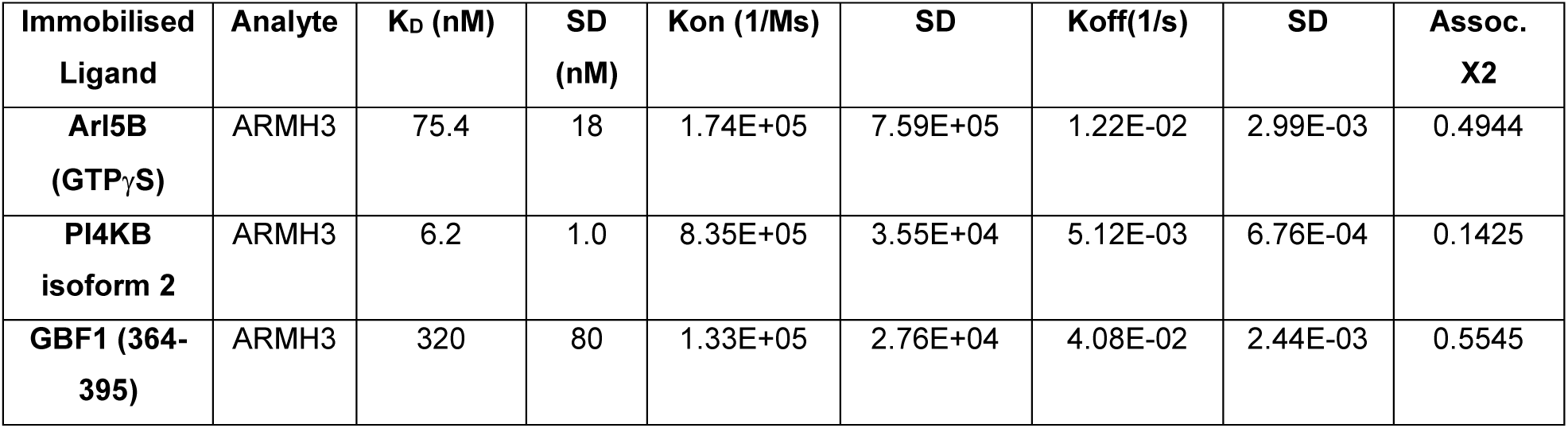
Binding constants for the different ARMH3 complexes.

**Table S2.**
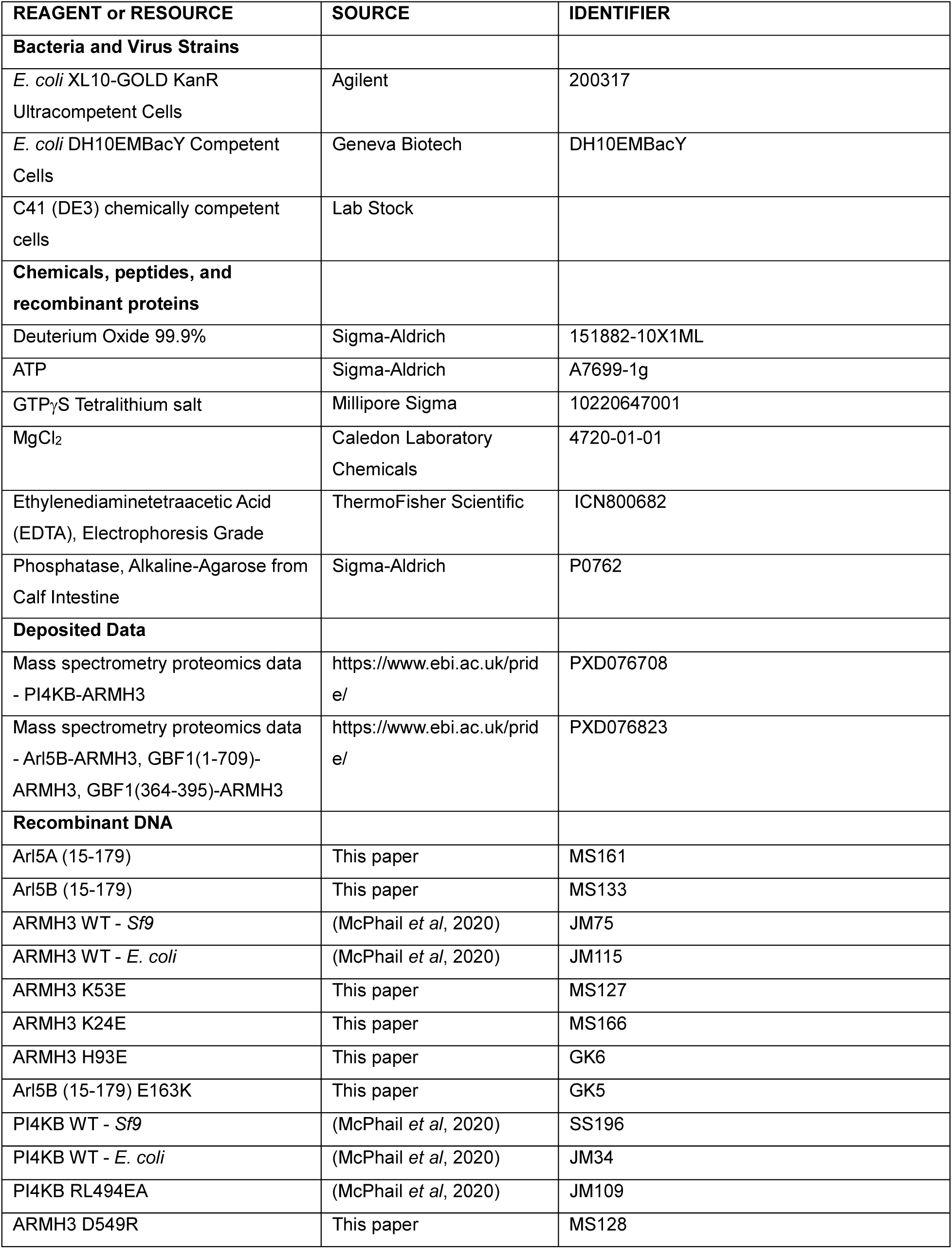
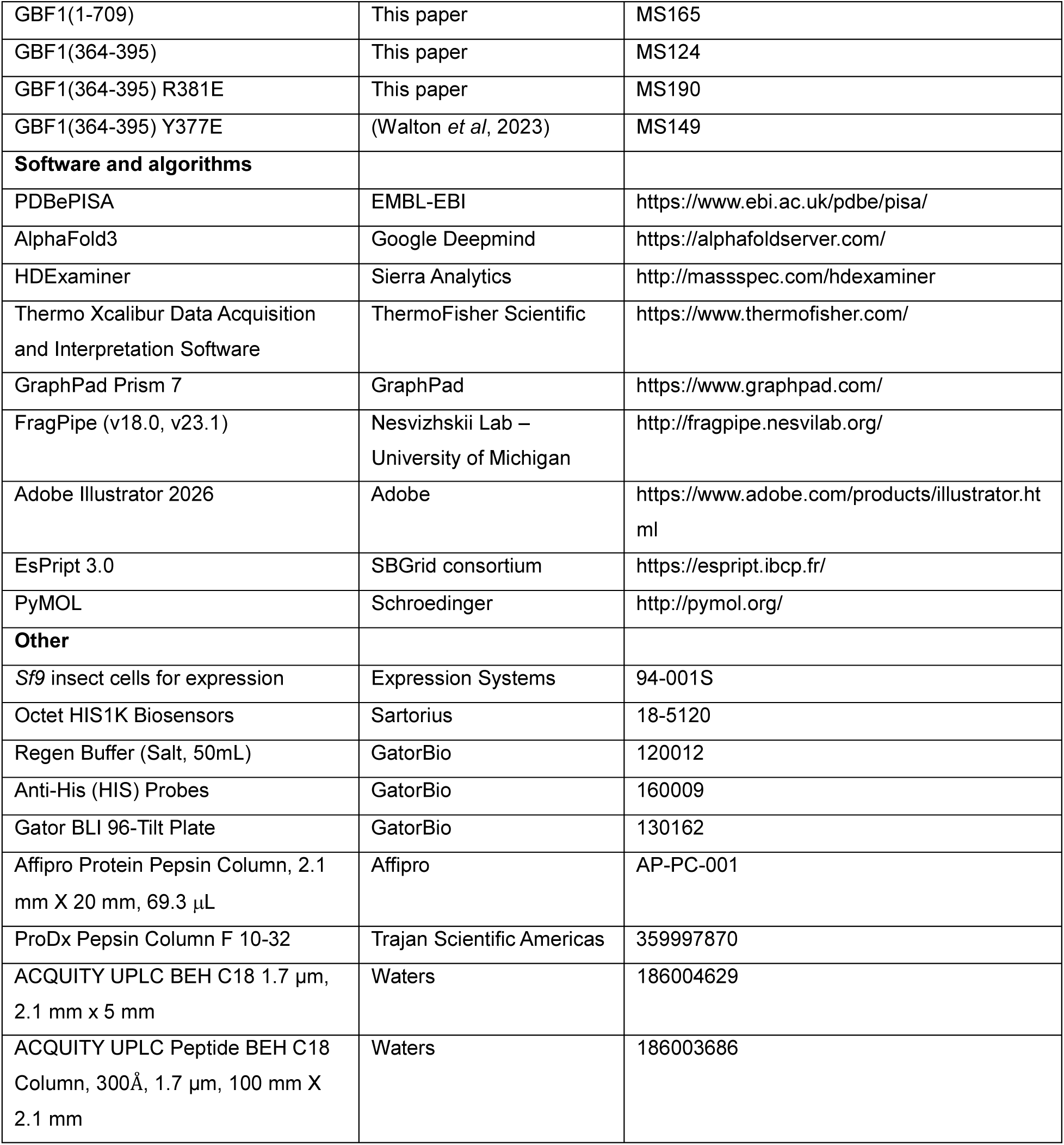
Key reagents/resources.

